# c-di-AMP is essential for the virulence of *Enterococcus faecalis*

**DOI:** 10.1101/2021.05.19.444760

**Authors:** Shivani Kundra, Ling Ning Lam, Jessica K. Kajfasz, Leila G. Casella, Marissa J. Andersen, Jacqueline Abranches, Ana L. Flores-Mireles, José A. Lemos

## Abstract

Second messenger nucleotides are produced by bacteria in response to environmental stimuli and play a major role in the regulation of processes associated with bacterial fitness, including but not limited to osmoregulation, envelope homeostasis, central metabolism, and biofilm formation. In this study, we uncovered the biological significance of c-di-AMP in the opportunistic pathogen *Enterococcus faecalis* by isolating and characterizing strains lacking genes responsible for c-di-AMP synthesis (*cdaA*) and degradation (*dhhP* and *gdpP*). Using complementary approaches, we demonstrated that either complete loss of c-di-AMP (Δ*cdaA* strain) or c-di-AMP accumulation (Δ*dhhP*, Δ*gdpP* and Δ*dhhP*Δ*gdpP* strains) drastically impaired general cell fitness and virulence of *E. faecalis*. In particular, the Δ*cdaA* strain was highly sensitive to envelope-targeting antibiotics, was unable to multiply and quickly lost viability in human serum or urine *ex vivo*, and was avirulent in an invertebrate (*Galleria mellonella*) and in two catheter-associated mouse infection models that recapitulate key aspects of enterococcal infections in humans. In addition to evidence linking these phenotypes to altered activity of metabolite and peptide transporters and inability to maintain osmobalance, we found that the attenuated virulence of Δ*cdaA* could be also attributed to a defect in Ebp pilus production and activity that severely impaired biofilm formation under both *in vitro* and *in vivo* conditions. Collectively, these results reveal that c-di-AMP signaling is essential for *E. faecalis* pathogenesis and a desirable target for drug development.

**IMPORTANCE:** *Enterococcus faecalis* is an opportunistic pathogen and leading cause of multidrug resistant hospital-acquired infections. During the course of an infection, bacteria encounter multiple adverse (stress) conditions and understanding the adaptive mechanisms used by pathogens to survive these stresses can facilitate the development of new antimicrobial therapies. Here, we used *in vitro*, *ex vivo* and *in vivo* approaches to determine the importance of the second messenger nucleotide c-di-AMP, a global regulator essential for bacterial adaptation to osmotic stress, to *E. faecalis* pathophysiology. We demonstrated that either accumulation of c-di-AMP or complete loss of c-di-AMP impaired cell fitness and virulence of *E. faecalis*. Remarkably, the strain that was unable to produce c-di-AMP was avirulent in three animal infection models indicating that c-di-AMP signaling is essential for *E. faecalis* pathogenesis and a suitable antimicrobial target.

## INTRODUCTION

Second messenger nucleotides are synthesized by bacteria in response to internal or external stimuli and are used to reprogram cell physiology through physical interactions with proteins (allosteric regulation), RNA riboswitches, or both (1-3). Despite being a relatively recent discovery, cyclic di-adenosine monophosphate (c-di-AMP) has been shown to control a variety of bacterial processes including osmoregulation, cell envelope homeostasis, stress and antibiotic tolerance, biofilm formation, central metabolism, DNA repair, genetic competence and sporulation (4-16). In addition, c-di-AMP plays an important role in host-pathogen interactions as it can be exported to the bacterial extracellular milieu promoting a potent type I interferon immune response (17). Two classes of enzymes have been identified as controlling c-di-AMP metabolism; diadenylate cyclases (DAC) are responsible for c-di-AMP synthesis (from two molecules of ATP or ADP), while phosphodiesterases (PDE) degrade c-di-AMP into 5’-Phosphoadenylyl-(3’ −> 5’)-adenosine (5’-pApA) and/or AMP (18). Most bacteria possess a single DAC with spore-forming Bacilli and Clostridia being among the few exceptions encoding three and two DACs, respectively (18). Most bacteria encode at least two types of c-di-AMP PDEs, which have either DHH/DHHA1 (aspartate-histidine-histidine) or HD (histidine-aspartate) catalytic domains (18). These DAC and PDE enzymes are ubiquitous in Gram-positive bacteria but can also be found in Gram-negative bacteria and Archaea (15, 16). Initially, c-di-AMP was thought to be essential as DAC gene deletion strains were not viable, but subsequent studies discovered that c-di-AMP is dispensable if cells are grown in minimal media (10, 19-21). Moreover, intracellular accumulation of c-di-AMP, as in the case of PDE-null mutants or overactive DAC, was also detrimental to cell homeostasis. Thus, c-di-AMP is both essential and toxic and often referred to as an “essential poison” (6). As a result, a growing number of studies have shown that disruption of c-di-AMP homeostasis can greatly diminish the virulence potential of bacterial pathogens (10, 11, 13, 22-24).

A natural resident of the human gastrointestinal tract, *Enterococcus faecalis* is also an opportunistic pathogen and leading agent of healthcare-associated infections, including wound infections, infective endocarditis, catheter-associated urinary tract infection (CAUTI) and central line-associated bloodstream infections (CLABSI) (25, 26). The inherent tolerance of *E. faecalis* to environmental stresses, including resistance to antibiotics, coupled with the abilities to develop robust biofilms on indwelling medical devices and host tissues and to subvert the immune system make enterococcal infections commonplace and difficult to eradicate (25, 27-30). Previous studies from our group demonstrated that another regulatory nucleotide, the stringent response effector (p)ppGpp, like c-di-AMP, regulates central metabolism, stress tolerance and biofilm formation in *E. faecalis*, and that complete lack of (p)ppGpp greatly increases antibiotic sensitivity and attenuates virulence of *E. faecalis* (31-34). Of interest, evidence indicates that the (p)ppGpp and c-di-AMP signaling networks are intertwined in members of the Firmicutes phylum. Specifically, the activity of GdpP and of PgpH is inhibited by (p)ppGpp, such that accumulation of (p)ppGpp leads to c-di-AMP accumulation (35, 36). At the same time, two recent studies showed that a previously uncharacterized c-di-AMP-binding protein (named CbpB in *L. monocytogenes* and DarB in *B. subtilis*) is a direct activator of the bifunctional (p)ppGpp synthetase/hydrolase Rel enzyme (also known as Rsh or RelA), and that c-di-AMP specifically binds to CbpB/DarB to inhibit its interaction with Rel and, consequently, impairs (p)ppGpp synthesis (37, 38). In addition, the inability of an *L. monocytogenes* DAC mutant to grow in complex media was overcome by inactivation of all of its (p)ppGpp synthetases thereby linking (p)ppGpp toxicity to c-di-AMP essentiality (19).

In this study, we sought to define the importance of c-di-AMP signaling to *E. faecalis* pathophysiology. We identified a single DAC (*cdaA*) and two PDE enzymes (*dhhP* and *gdpP*) and showed that genetic manipulations of these genes that lead to either a complete loss of c-di-AMP (Δ*cdaA*) or c-di-AMP accumulation (Δ*dhhP*, Δ*gdpP* and Δ*dhhP*Δ*gdpP*) have profound consequences to the cell fitness and virulence potential of *E. faecalis*. Most notably, we showed that c-di-AMP is absolutely critical to *E. faecalis* pathogenesis as the Δ*cdaA* strain was avirulent in multiple animal infection models. In addition to evidence linking c-di-AMP dysregulation to defects in metabolite and peptide transport and the inability to maintain osmotic balance, we found that the attenuated virulence of Δ*cdaA* could be also attributed to a defect in Ebp pilus production and activity, which severely impaired biofilm formation *in vitro* and *in vivo*. Collectively, our results underpin that antimicrobial therapies targeting c-di-AMP enzymes and, perhaps, c-di-AMP effector molecules can facilitate the development of new antimicrobial therapies.

## RESULTS

### Identification of the enzymes responsible for synthesis and degradation of c-di-AMP in *E. faecalis*

Through bioinformatic analysis using the amino acid sequences of DAC and PDE orthologues from closely-related streptococci, we identified one DAC-encoding gene (gene ID. *OG1RF_RS08715*), and two PDE-encoding genes (*OG1RF_RS06025* and *OG1RF_RS00060*) in the *E. faecalis* OG1RF genome; all three genes are part of the *E. faecalis* core genome. Similar to CdaA orthologues (18), the *E. faecalis* DAC displays a C-terminal DAC domain and three transmembrane helices at the N-terminal domain. Both PDE proteins contain conserved DHH-DHHA1 catalytic domains and, while *OG1RF_RS06025* is predicted to be a standalone DHH-DHHA1 cytoplasmic protein, *OG1RF_RS00060* possess two N-terminal transmembrane domains, followed by a PAS (Per-Arnt-Sim) domain, a degenerate GGDEF domain and the N-terminally located DHH-DHHA1 domain. Based on the presence of these conserved domains and high similarity with previously characterized enzymes, we adopted the *cdaA* (*OG1RF_RS08715*), *dhhP* (*OG1RF_RS06025*) and *gdpP* (*OG1RF_RS00060*) nomenclature to rename these genes. Here, we note that the *dhhP* gene is also known as *pde* or *pde2* in closely-related bacteria (11, 39).

To confirm the suspected functions of the *cdaA*, *dhhP* and *gdpP* gene products, single (Δ*cdaA*, Δ*dhhP*, Δ*gdpP*) and double (Δ*dhhP*Δ*gdpP*) mutants were generated using a markerless in-frame deletion strategy (40). Based on the expectation that c-di-AMP is essential for growth in complex media, the Δ*cdaA* strain was isolated in a chemically-defined media (CDM) supplemented with 20 mM glucose (41). Upon confirmation that the mutations occurred as designed by Sanger sequencing, we used LC-MS/MS to determine the intracellular levels of c-di-AMP in exponentially-grown CDM cultures of parent (OG1RF) and mutant (Δ*cdaA*, Δ*dhhP*, Δ*gdpP* and Δ*dhhP*Δ*gdpP*) strains (Fig. 1A). In agreement with the predicted fucntion, no detectable c-di-AMP was found in the Δ*cdaA* strain (herein also referred as c-di-AMP^0^ strain) revealing that CdaA is the only enzymatic source of c-di-AMP in *E. faecalis*. In addition, high intracellular c-di-AMP pools were detected in the Δ*gdpP* (∼3-fold) and Δ*dhhP* (∼5-fold) mutants when compared to the parent strain. The increase in c-di-AMP in single PDE mutants was additive, as the Δ*dhhP*Δ*gdpP* double mutant displayed ∼8-fold increase in c-di-AMP pools. Because suppressor mutations are known to arise in DAC mutants, we subjected the Δ*cdaA* isolate to whole-genome sequencing (WGS) analysis, which ruled out the presence of additional mutations.

**FIG 1.**
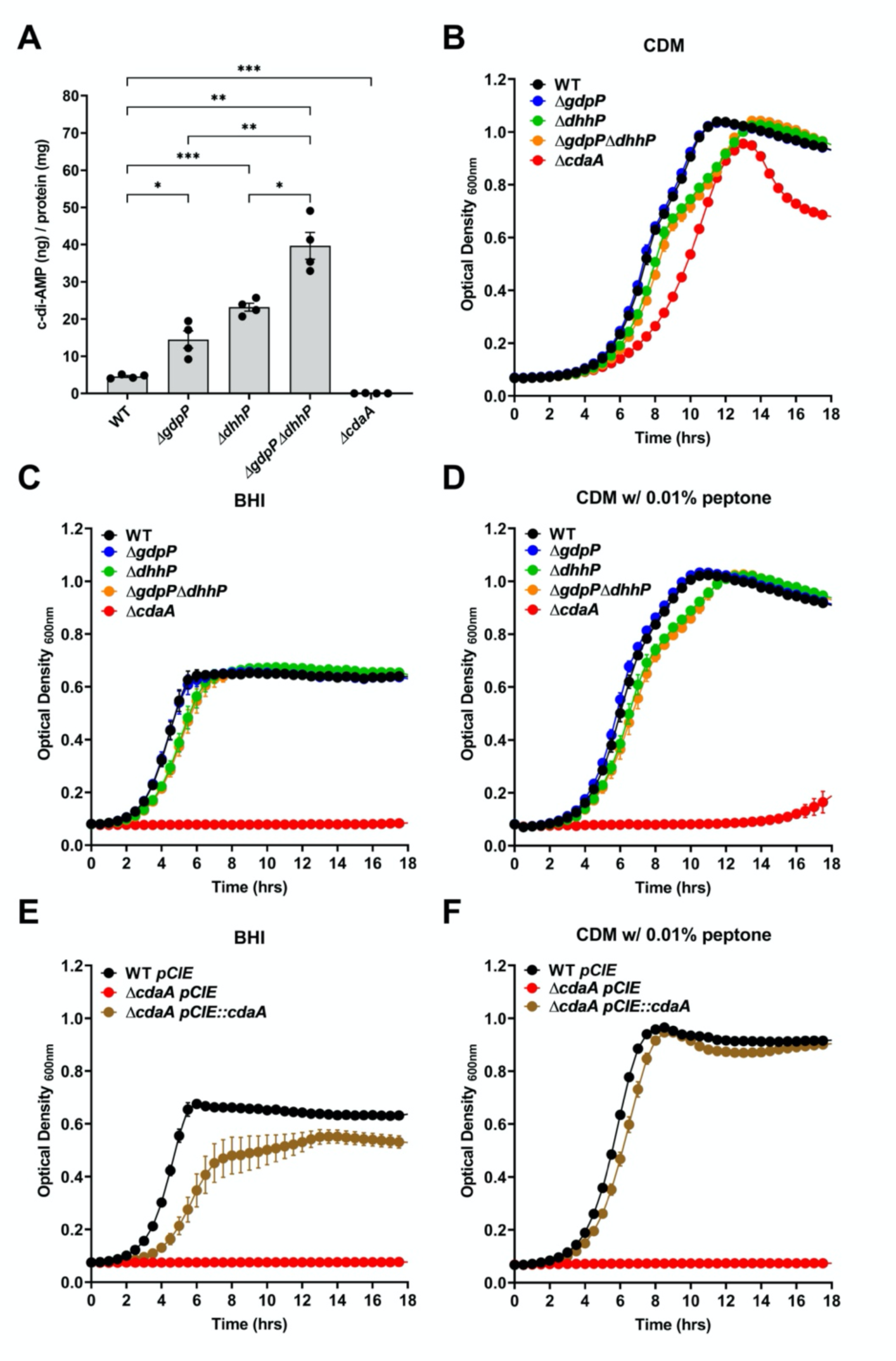
Growth characteristics of DAC and PDE mutants. (A) LC-MS/MS quantification of c-di-AMP in *E. faecalis* OG1RF (WT) and indicated mutant strains grown to mid-log in CDM. Statistical analysis was performed using unpaired *t*-test with Welch’s correction on four biological replicates. Error bars represent the standard error of margin (SEM). * *p* ≤ 0.05, ** *p* ≤ 0.01 and *** *p* ≤ 0.001. (B-F) Growth curves of *E. faecalis* WT and mutants in CDM (B), BHI (C), CDM with 0.01% peptone (D), and genetic complementation of Δ*cdaA* in BHI (E) and CDM + 0.1% peptone (F). Data points represent the average and error bars represent the standard error of margin of nine biological replicates.

### c-di-AMP is essential for growth in the presence of peptides and osmolytes

The essentiality of c-di-AMP for growth in complex media or in minimal media supplemented with osmolytes or peptides has been demonstrated in several bacteria, including *B. subtilis* (20), *L. monocytogenes* (19) and *S. aureus* (21). Thus, we determined the ability of the Δ*cdaA*, Δ*dhhP*, Δ*gdpP* and Δ*dhhP*Δ*gdpP* strains to grow in the complex media BHI or in CDM supplemented with low molecular weight (MW) osmolytes or peptides. In low salt and peptide free CDM, the Δ*cdaA* strain grew slower with a slightly extended lag phase and lower growth yields (Fig. 1B). However, Δ*cdaA* failed to grow in BHI or CDM supplemented with as little as 0.01% peptone (Fig 1C-D). Under these conditions, the Δ*dhhP* and Δ*dhhP*Δ*gdpP* strains showed modest growth defects whereas the Δ*gdpP* strain phenocopied the OG1RF parent strain (Fig 1B-D). We also tested the ability of the mutants to grow in CDM supplemented with the low MW osmolytes carnitine and glycine betaine. In both cases, the addition of either one of the osmolytes abolished growth of Δ*cdaA* while not having an impact on growth of parent or any PDE mutant strain (Fig S1). Genetic (*in*-*trans*) complementation rescued the growth defect of Δ*cdaA* in BHI as well as in CDM supplemented with peptone, glycine betaine or carnitine (Fig 1E-F and Fig S1). Next, we took advantage of the stable nature of cyclic nucleotides to ask if exogenous addition of purified c-di-AMP restored growth defects of the Δ*cdaA* strain. Remarkably, purified c-di-AMP restored the ability of Δ*cdaA* to grow in CDM supplemented with 1% peptone in a dose-dependent manner (Fig S2A). Interestingly, exogenous c-di-AMP did not rescue the growth of Δ*cdaA* in BHI (Fig S2B), even if supplied at higher concentrations (up to 25 μM c-di-AMP, data not shown).

In most bacteria studied to date, the essentiality of c-di-AMP is directly associated to its role in K^+^ homeostasis (20, 42, 43). However, the effect of K^+^ on growth of c-di-AMP mutants varies according to the composition of the growth media and bacterial species. In *B. subtilis*, a strain lacking c-di-AMP is fully viable under low K^+^ conditions but unable to survive high K^+^ concentrations (20). Paradoxically, increasing osmotic pressure with addition of salts (KCl or NaCl) rescued growth defects of *L. monocytogenes* and *S. aureus* c-di-AMP^0^ strains in complex media (12, 21). Similar to the latter, we found that addition of high slat comncentrations (250 to 500 mM KCl or NaCl) partially restored growth of Δ*cdaA* in CDM-peptone, even though high salt concentrations were mildly inhibitory to cell growth (Fig S3A-D). To determine the salt tolerance of Δ*cdaA*, we compared the ability of OG1RF and Δ*cdaA* strains to grow in (peptone-free) CDM supplemented with 250 or 500 mM of either KCl or NaCl. Interestingly, omission of peptone from the growth media enhanced salt sensitivity of both parent and mutant strains indicating that the protein/peptide/amino acid mixture counteracted the negative effect of salt stress (Fig. S3).

### c-di-AMP contributes to *E. faecalis* antibiotic tolerance

In other bacteria, alterations in c-di-AMP concentrations lead to defects in cell division, chain length, aberrant morphology and increased sensitivity to cell envelope-targeting antibiotics (10, 13, 44-46). Using transmission electron microscopy (TEM) and scanning electron microscopy (SEM), we observed an aberrant morphology for the Δ*cdaA* strain displaying enlarged cells with irregular division patterns and a high number of lysed/dead cells (Fig. S4); however, the morphologic characteristics of the Δ*dhhP*Δ*gdpP* strain were unremarkable, largely resembling the parent strain. Next, we tested the susceptibility of DAC and PDE mutants to antibiotics that target cell wall biosynthesis (ampicillin, bacitracin and vancomycin) or the cell membrane (daptomycin). In general, the Δ*dhhP*, Δ*dhhP*Δ*gdpP* and Δ*cdaA* strains showed increased sensitivity to all four antibiotics albeit the differences observed in the presence of ampicillin or vancomycin were not significant or were significant though modest (Fig. 2A-B). On the other hand, the Δ*dhhP*, Δ*dhhP*Δ*gdpP* and Δ*cdaA* strains were highly susceptible to bacitracin or daptomycin (Fig. 2C-D). Once again, the Δ*gdpP* strain phenocopyed the parent strain.

**FIG 2.**
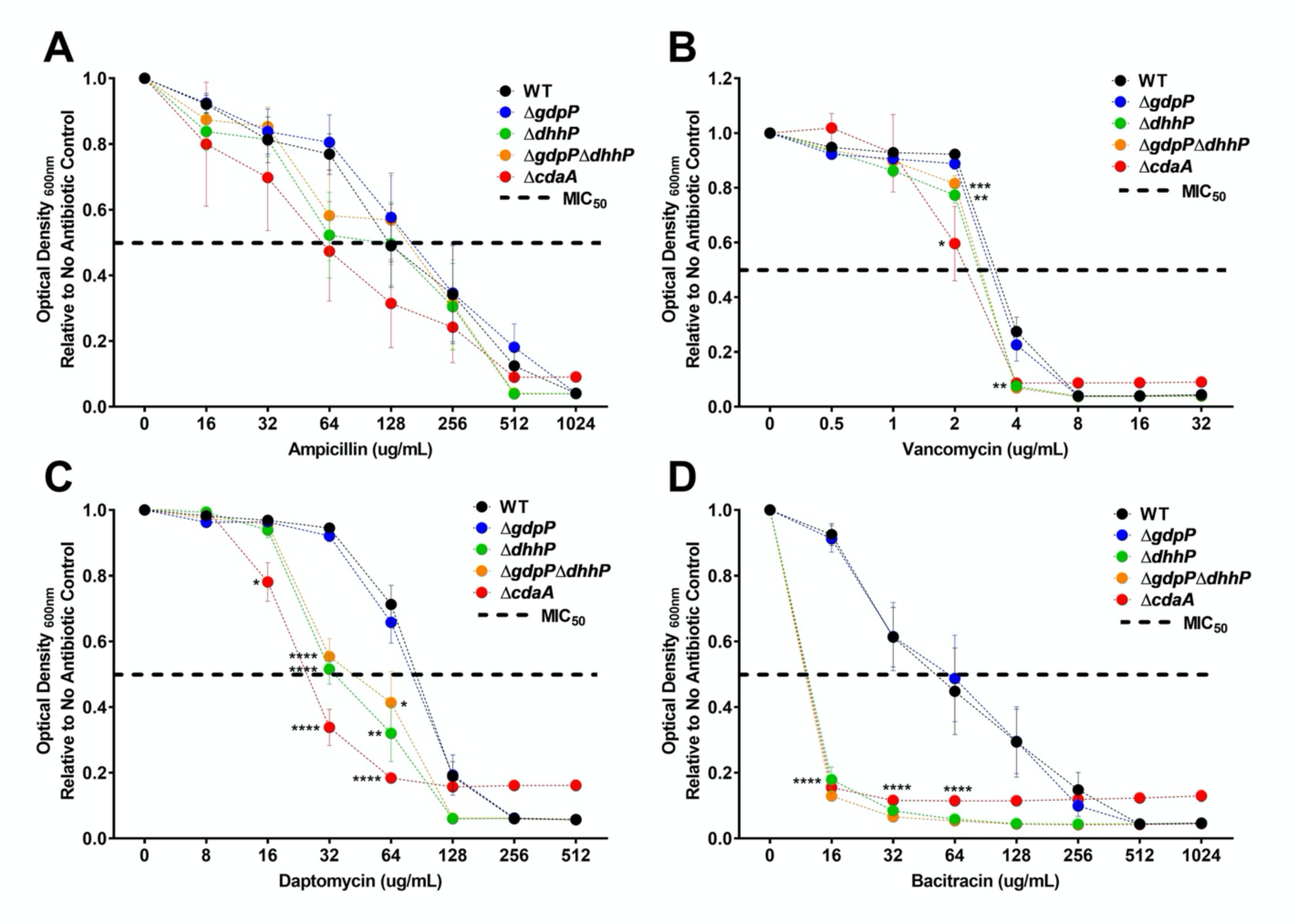
Antibiotic susceptibility of DAC and PDE mutants. Final growth yields of *E. faecalis* OG1RF (WT) and indicated mutants after 24 hr incubation in CDM supplemented with 2-fold increasing concentrations of ampicillin (A), vancomycin (B), daptomycin(C) and bacitracin (D). Data points represent the average of nine biological replicates. Error bars represent the standard error of margin (SEM). Statistical analysis was performed using unpaired *t*-test with Welch’s correction. Error bar represents the standard error of margin. * *p* ≤ 0.05, ** *p* ≤ 0.01, *** *p* ≤ 0.001, **** *p* ≤ 0.0001.

### c-di-AMP mediates growth and survival in human serum and urine

Next, we assessed the ability of parent and mutant strains to grow and survive in pooled human serum or urine *ex vivo*. The PDE mutants grew and remained fully viable for 24 h in either serum or urine with modest differences in growth yields observed for Δ*dhhA* and Δ*dhhA*Δ*gdpP* in urine. Importantly, the Δ*cdaA* strain was unable to grow and quickly lost its viability with ∼1-log CFU reduction in serum and ∼3.5-log CFU reduction in urine after 8-h of incubation (Fig. 3A-B). The growth/survival defect of Δ*cdaA* was fully restored in the genetically-complemented strain (Fig. 3C-D). Of interest, we recently showed that transcription of *dhhP* and *gdpP* was strongly induced when a mid-log grown culture of *E. faecalis* OG1RF was transferred to human urine (32). Here, we found that transcription of the *gdpP* and *dhhP* genes was also induced after a 30 min incubation cells in human serum (∼2.5 and 6-fold, respectively) whereas *cdaA* mRNA was repressed by ∼2-fold (Fig. 3E). In concurrence with the transcriptional patterns, we detected ∼6-fold reduction in c-di-AMP when comparing intracellular c-di-AMP pools of exponentially-growing cell in CDM to cells incubated in serum or urine for 30 minutes (Fig. 3F).

**FIG 3.**
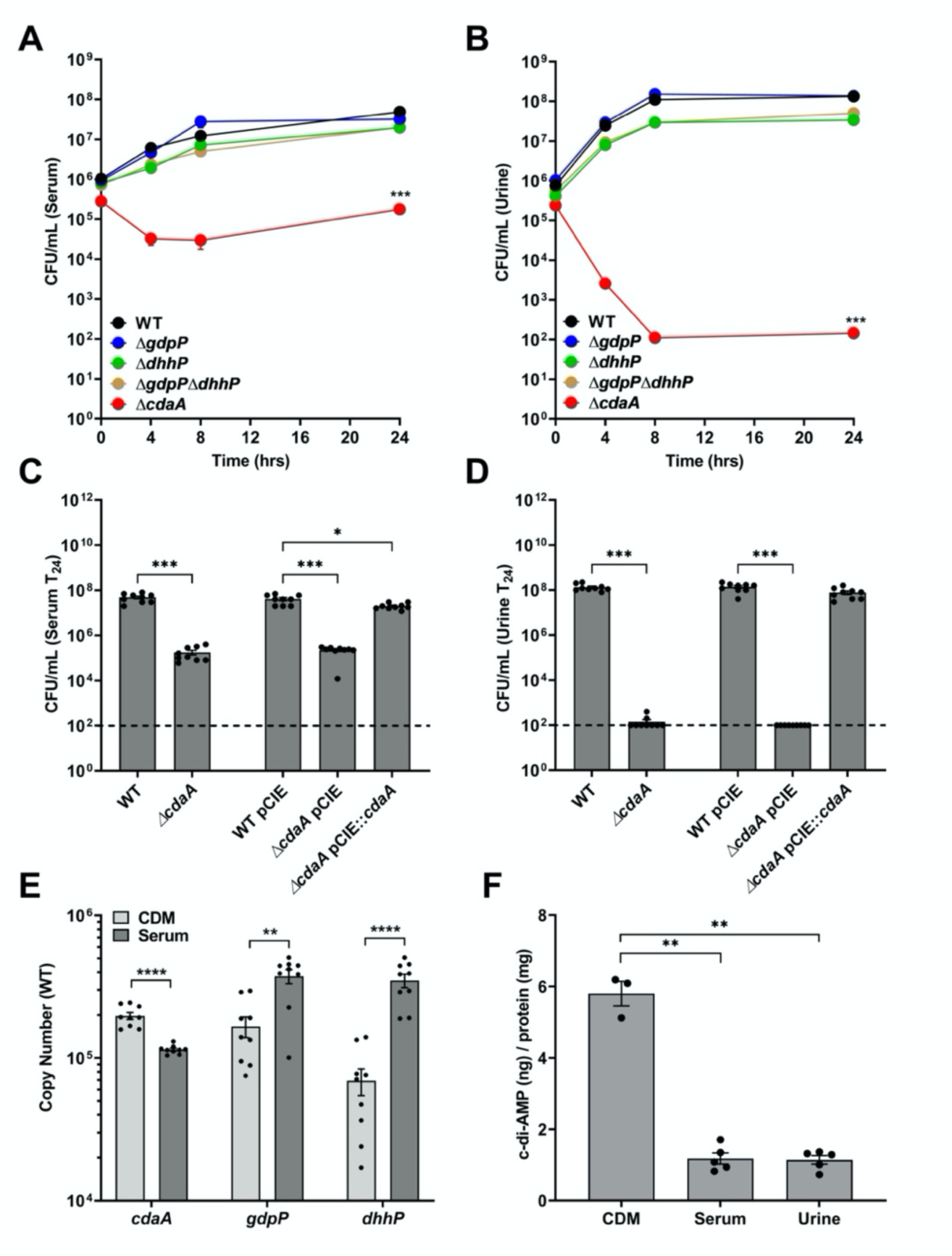
Growth and survival of DAC and PDE mutants in serum or urine. (A-B) Colony-forming units (CFU) of *E. faecalis* OG1RF (WT) and indicated mutants incubated in pooled human serum (A), or pooled human urine (B). (C-D) CFU counts of WT, Δ*cdaA* and complemented Δ*cdaA* (Δ*cdaA* pCIE::*cdaA*) after 24 hrs incubation in pooled serum (C) or pooled urine (D). Data points represent average of nine biological replicates and error bars represent the standard error of margin. (E) Comparison of reversed transcribed cDNA copy number of *cdaA*, *gdpP* and *dhhP* in *E. faecalis* OG1RF grown to mid-log phase in CDM and 30 min after transfer to human serum. (F) LC-MS/MS quantification of c-di-AMP in *E. faecalis* OG1RF (WT) grown to mid-log phase in CDM or 30 min after transfer to serum or urine. Data points in panel E and F represent average of nine and five biological replicates, respectively. Dashed lines in panels C and D indicate limit of detection. Statistical analysis was performed using one-way ANOVA with Dunnett’s multiple comparison test. * *p* ≤ 0.05, ** *p* ≤ 0.01, *** *p* ≤ 0.001, **** *p* ≤ 0.0001.

### c-di-AMP is critical for *E. faecalis* virulence in multiple animal infection models

Presently, a direct assessment of the significance of c-di-AMP in bacterial pathogenesis is limited to a handful of bacteria, with most studies limited to strains that accumulate c-di-AMP (10, 11, 45, 47, 48). Here, we used three well-established animal infection models to uncover the significance of c-di-AMP to *E. faecalis* pathogenesis. In the *Galleria mellonella* invertebrate model (Fig. 4A), virulence of Δ*dhhP* and Δ*dhhP*Δ*gdpP* strains was attenuated with Δ*gdpP* showing a similar but not statistically significant trend. Most notably, the Δ*cdaA* strain was nonpathogenic to *G. mellonella* (Fig. 4A). Next, we used two foreign body-associated mouse infection models that recapitulate some of the environmental and immunological conditions that promote enterococcal infections in humans. In a catheter-associated peritonitis mouse model (49), all three mutants tested (Δ*cdaA*, Δ*dhhP* and Δ*dhhP*Δ*gdpP*) were recovered in significantly lower numbers from peritoneal washes, implants or spleens (Fig. 4B). The total bacteria recovered from animals infected with Δ*dhhP* and Δ*dhhP*Δ*gdpP* was nearly identical (∼1 log10 CFU reduction) and, as expected, the most dramatic differences were observed in animals infected with Δ*cdaA* (≥2 log10 CFU reduction), with no Δ*cdaA* colonies isolated from implanted catheters (Fig. 4B). Following the exact same pattern, the Δ*cdaA* strain was virtually nonpathogenic in a CAUTI mouse model whereas Δ*dhhP* and, even more so, Δ*dhhP*Δ*gdpP* strains showed intermediate phenotypes (Fig. 4C).

**FIG 4.**
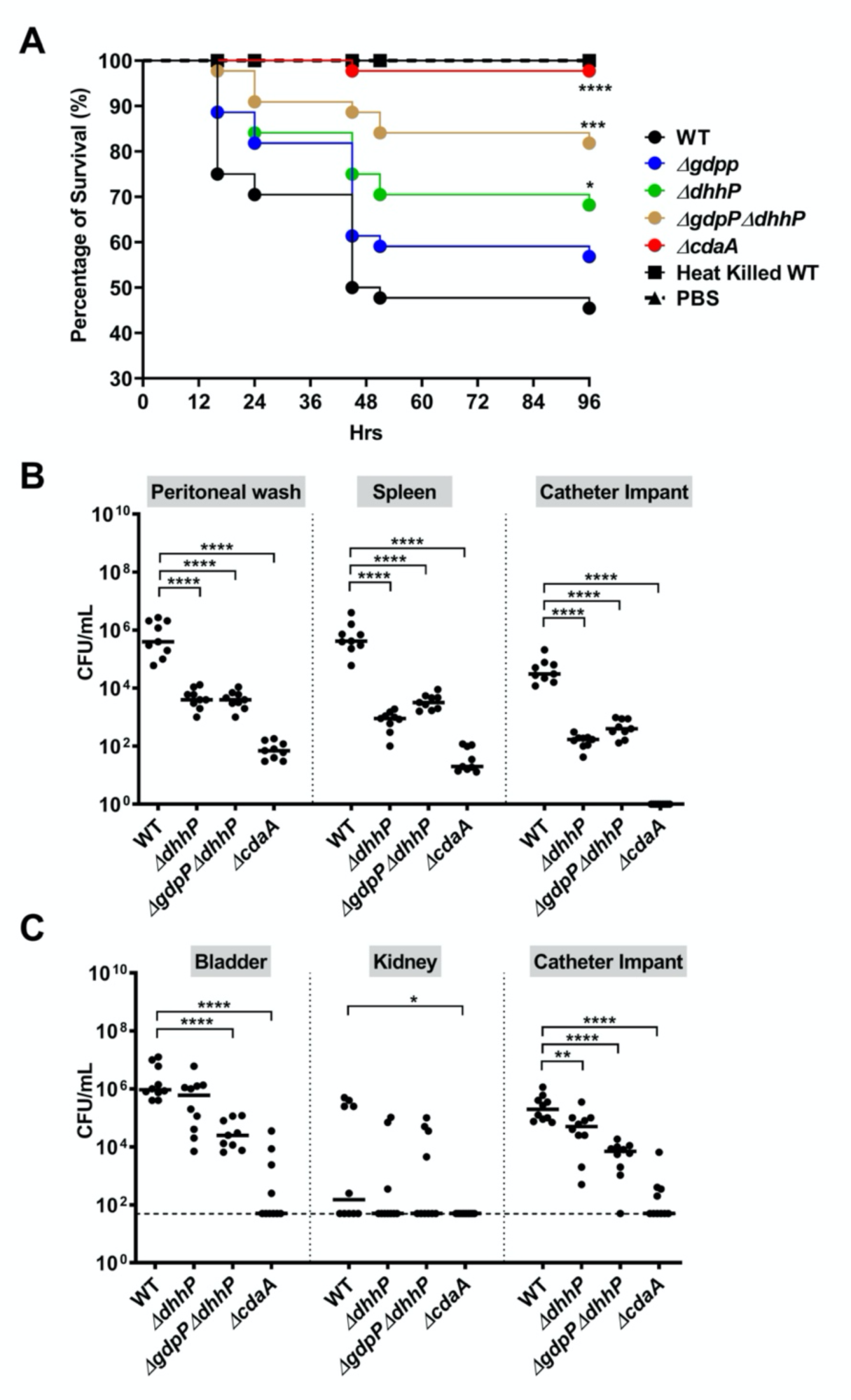
Virulence potential of DAC and PDE mutants in different animal models. (A) Percentage survival of *G. mellonella* 96 hrs post-infection with *E. faecalis* OG1RF (WT) or the indicated mutants. Each curve represents a group of 15 larvae injected with ∼1 x 10^5^ CFU of *E. faecalis*. Data points represent average of 3 biological replicates. Statistical analysis was performed using Log-rank (Mantel-Cox) test. (B) Total CFU recovered after 48 hrs from peritoneal wash, spleen or catheter of mice infected with 2 x 10^8^ CFU of WT or the indicated mutants. (C) Total CFU recovered after 24 hrs from bladder, kidney or catheter of mice infected with 1 x10^7^ CFU of WT or the indicated mutants. For (B-C), data points represent nine mice infected with three biological replicates. Bars represent the median. Statistical analysis was performed using Mann-Whitney test. * *p* ≤ 0.05, ** *p* ≤ 0.01, *** *p* ≤ 0.001, **** *p* ≤ 0.0001. The dashed line represents the limit of detection.

### c-di-AMP mediates biofilm formation in an Ebp-dependent manner

In both mouse models, the virtual inability of the Δ*cdaA* strain to colonize the catheters indicates that c-di-AMP modulates expression and/or activity of virulence factors associated with biofilm formation.

Using a microtiter plate-based assay, we confirmed this prediction as the Δ*cdaA* strain formed very poor biofilms after 18 hrs of incubation (Fig. 5A). In *E. faecalis*, robust biofilm formation on host tissue surfaces and urinary catheters is directly associated with expression of the EbpABC pilus; strains lacking *ebp* genes form poor biofilms on different types of surfaces, in particular fibrinogen-coated surfaces, and display attenuated virulence in CAUTI and infective endocarditis models (50, 51). To evaluate if the *in vitro* and *in vivo* biofilm defects of Δ*cdaA* was linked to the Ebp pilus, we first performed a 4-h *in vitro* adhesion assay incubating mid-log grown cells in CDM supplemented with 50 mM Na_2_CO_3_, a condition that stimulates Ebp synthesis (52). As expected, Na_2_CO_3_ increased surface adhesion by the parent strain but, most importantly, we found that the ability of Δ*cdaA* to adhere to the plate surface was further reduced under the Ebp-inducive condition (Fig. 5B). Moreover, inactivation of *dhhP*, *gdpP* or both did not affect *E. faecalis* biofilm formation or surface adhesion while genetic complementation of Δ*cdaA* rescued both phenotypes (Fig. 5A-B).

**FIG 5.**
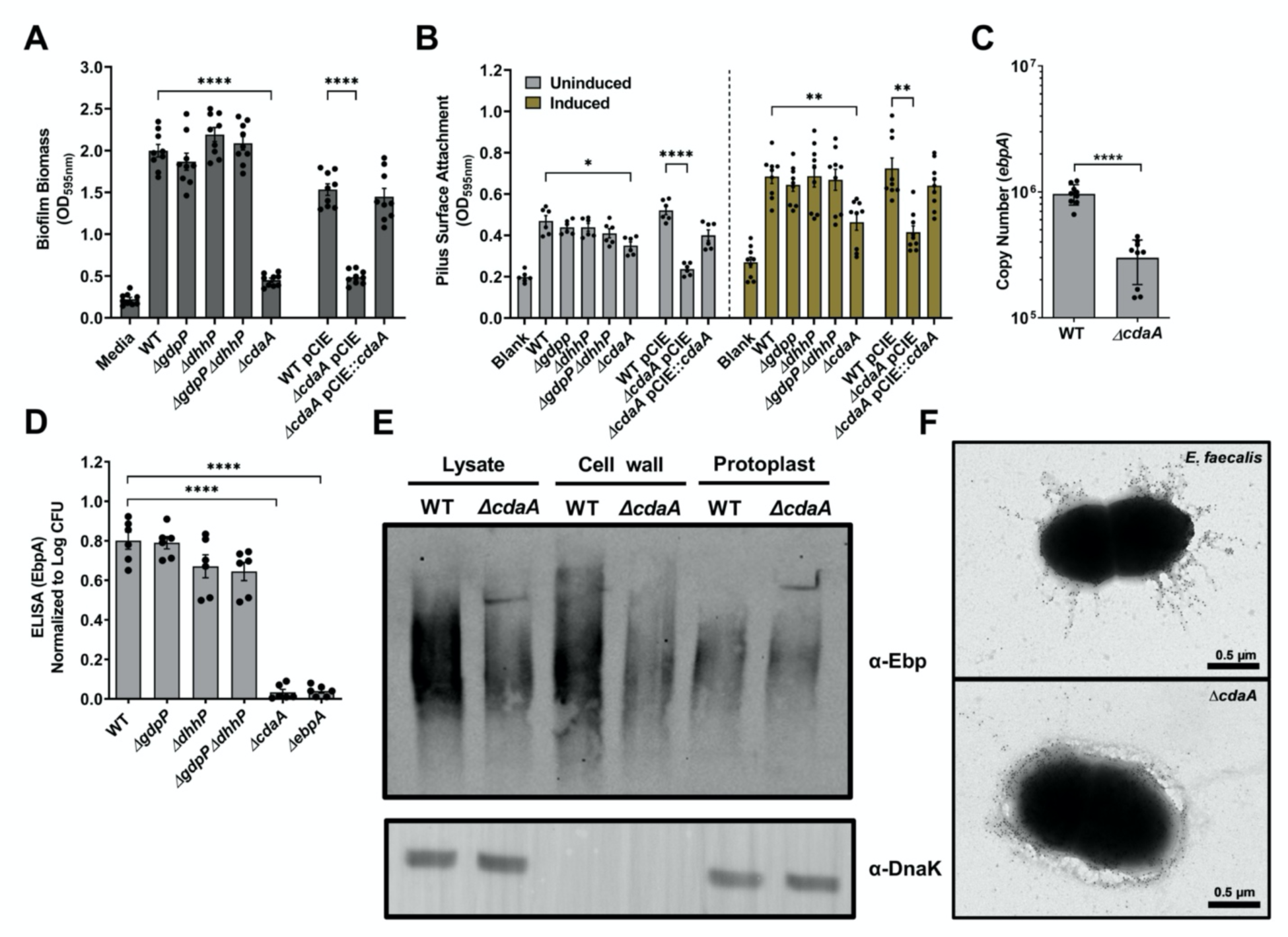
c-di-AMP mediates biofilm formation by modulating Ebp expression and biogenesis. (A) Adherence biofilm biomass quantification of *E. faecalis* OG1RF (WT) and indicated mutants grown in CDM for 24 hrs. (B) Adherence of WT and indicated mutants under Ebp-inducing condition (CDM + 50 mM sodium bicarbonate). Data points represent nine biological replicates assessed in three independent experiments. For (A-B), statistical analysis was performed using one-way ANOVA with Welch’s correction. (C) Comparison of reversed transcribed cDNA copy number of *ebpA* in *E. faecalis* WT and Δ*cdaA* strains grown to mid-log phase in CDM. Data points represent six biological replicates and statistical analysis was performed using unpaired *t*-test with Welch’s correction. ** *p* ≤ 0.01, *** *p* ≤ 0.001, **** *p* ≤ 0.0001. Error bars represent the standard error of margin. (D) Adherence ability of WT and indicated mutants via EbpA binding to EbpA-specific antibody using ELISA. Data points represent six biological replicates and statistical analysis was performed using one-way ANOVA with Dunnett’s multiple comparison test. **** *p* ≤ 0.0001. (E) Immunoblot showing Ebp expression in whole cell lysates, cell wall fraction and protoplasts of *E. faecalis* WT and Δ*cdaA* cells harvested during mid-log growth phase. Immunoblot using a polyclonal anti-DnaK antibody was used as protein loading control. The images shown are representative of three experiments using three different biological replicates. (F) Immunoelectron microscopy of WT and Δ*cdaA* strains using EbpA-specific antibody (1:1000).

Through qRT-PCR and ELISA, we found that *ebpA* transcription was significantly reduced (∼5-fold) in the Δ*cdaA* strain and that EbpA was below the detection limit in the ELISA, mirrowing the the Δ*ebpA* strain (Fig. 5C-D). However, immunoblot analysis revealed that EbpA was produced in the Δ*cdaA* strain, albeit in lower amounts when compared to OG1RF (Fig. 5E). Here, we speculated that the ELISA result was caused by the combination of reduced Ebp production and impaired function, possibly due to defects in protein translocation, pilus assembly or surface anchoring. To explore some of these possibilities, we used high-resolution TEM to visualize the spatial organization and abundance of the Ebp pilus on the cell surface. As shown in previous studies (53, 54), the Ebp pilus was detected as extracellular filamentous fibers extending outwards from the cell surface of OG1RF (Fig. 5F, top panel). However, these fiber-like structures were almost completely absent in Δ*cdaA*, with EbpA appearing more evenly distributed around the cell surface (Fig. 5F).

### The impact of c-di-AMP on global gene expression

To evaluate the impact of c-di-AMP dysregulation at a global level, we used RNA sequencing (RNA-seq) to compare the transcriptome of OG1RF, Δ*cdaA*, and Δ*dhhP*Δ*gdpP*. The complete absence of c-di-AMP (Δ*cdaA*) resulted in the differential expression of 302 genes (2-fold cutoff, 147 upregulated and 155 downregulated – Table S1), and c-di-AMP accumulation (Δ*dhhP*Δ*gdpP*) resulted in 163 differentially expressed genes (2-fold cutoff, 62 upregulated and 101 downregulated – Table S2). Notably, the *ebpABC* operon was among the most strongly repressed genes (4.5- to 11-fold downregulation) in Δ*cdaA* validating the qRT-PCR quantifications. Among the differently expressed genes, 77 were common for both mutants with 12 genes showing opposite pattern of expression. Among the genes with opposing expression patterns were genes from a transcriptional unit coding for a putative compatible solute ABC-type transporter (*OG1RF_RS10300-RS10305*) that was downregulated in Δ*cdaA* and upregulated in Δ*dhhP*Δ*gdpP*. Given the strong association of c-di-AMP with osmoregulation, it was interesting to also observe that transcriptional units coding for polyamine/osmolyte transporters (*OG1RF_RS10340-RS10355* and *OG1RF_RS05150-RS05165*) were downregulated in the Δ*cdaA* strain. Conversely, a glutamate transport operon (*OG1RF_RS06355-65*) and an annotated oligopeptide transporter (*OG1RF_RS00290*) were among the most highly upregulated genes in the Δ*cdaA* strains. Among genes showing the same expression trend in Δ*cdaA* and Δ*dhhA*Δ*gdpP* were a large transcriptional unit (*OG1RF_RS07320-RS07370*) coding for genes involved in pyrimidine metabolism, a cluster of surface-anchored WxL domain proteins with unknown function (*OG1RF_RS02570-RS02595*), the murein hydrolase regulator *lrgA*-*lrgB* genes (*OG1RF_RS12565-RS12570*) and the arginine deiminase (ADI) operon (*OG1RF_RS490-RS515*).

**TABLE 1.** Bacterial strains used in this study.

| Strains | Relevant characteristics | Source/reference |
| --- | --- | --- |
| *E. faecalis* |  |  |
| OG1RF | Wild-type strain; Rif<sup>r</sup> Fus<sup>r</sup> | Laboratory stock |
| CK111 | OG1Sp *upp4::P<sub>23</sub>repA4*, Spec<sup>r</sup> | Laboratory stock |
| $\Delta gdpP$ | *gdpP* (OG1RF_RS00060) deletion | This study |
| $\Delta dhhP$ | *dhhP* (OG1RF_RS06025) deletion | This study |
| $\Delta gdpP\Delta dhhP$ | *gdpP* and *dhhP* double deletion | This study |
| $\Delta cdaA$ | *cdaA* (OG1RF_RS08715) deletion | This study |
| $\Delta cdaA$-pCIE::*cdaA* | genetically-complemented $\Delta cdaA$, Cm<sup>r</sup> | This study |
| $\Delta cdaA$-pCIE | $\Delta cdaA$ empty pCIE vector, Cm<sup>r</sup> | This study |
| $\Delta ebpA$ | *ebpA* (OG1RF_RS04550) deletion | (54) |
| *E. coli* |  |  |
| DH10b | Cloning host | Laboratory stock |
| EC1000 | Host for cloning RepA-dependent plasmids | Laboratory stock |

## DISCUSSION

Since its serendipitous discovery in 2008 (55), c-di-AMP quickly emerged as one of the most important second messenger nucleotides of bacteria that exerts critical regulatory roles in a number of cellular processes, including (but not restricted to) osmoregulation, cell envelope homeostasis, and biofilm formation (15, 16, 43, 56). Furthermore, accumulating evidence that c-di-AMP mediates bacterial virulence and recognition that the enzymes that synthesize and degrade c-di-AMP are absent in eukaryotes caught the attention of investigators that are actively pursuing the identification of c-di-AMP inhibitors for future clinical application. At first, c-di-AMP was thought to be essential as DAC-null strains did not grow in common laboratory media unless they acquired suppressor mutations (9-12, 19-21, 37, 57, 58). Later, it was discovered that c-di-AMP essentiality could be overcome by growing DAC mutants (c-di-AMP^0^ strains) in minimal media (19-21, 58). It followed that loss of osmobalance control caused by lack of c-di-AMP regulation, could be tolerated in media free of osmolytes and peptides, which in turn also provided an explanation for the emergence of suppressor DAC mutants with secondary mutations in genes coding for K^+^, peptide or osmolyte transporters. While a genetically-stable *E. faecalis* DAC mutant (Δ*cdaA*) was viable and grew relatively well in CDM, if challenged to grow in BHI or CDM supplemented with peptone, our Δ*cdaA* strain also acquired suppressor mutations that restored the capacity to grow in complex media (data not shown). WGS analysis of six suppressor mutant isolates identified SNPs in genes coding for transport proteins targeted by c-di-AMP in other bacteria, which included K^+^ and compatible solute ABC-type transporters.

However, it should be noted that the quasi-essentiality of c-di-AMP is not universal among Firmicutes, as a genetically-stable Δ*cdaA* strain that grew in complex media has been isolated and then characterized in *S. pyogenes* (10). Interestingly, we also found that exogenous addition of c-di-AMP reverts the growth defect phenotype of Δ*cdaA* in CDM supplement with peptone but not in BHI (Fig. S2). Because c-di-AMP is negatively charged and, in theory, should not diffuse through the cytoplasmic membrane, this result suggests that c-di-AMP can be actively imported, either via a promiscuous transporter or by a dedicated c-di-AMP transporter. This finding also suggests that metabolites present in BHI somehow inhibit c-di-AMP uptake. While several investigations have shown that bacteria respond to exogenous c-di-GMP, to our knowledge, only one previous report demonstrated the effect of exogenous c-di-AMP on expression of c-di-AMP-dependent phenotypes. Specifically, exogenous c-di-AMP, mixed with cationic polyamines to facilitate internalization, was shown to promote *B. subtillis* sporulation (14). With the understanding that working with physiologically-relevant will be crucial, exogenously added c-di-AMP can be used to further explore the scope of c-di-AMP regulation and to identify new c-di-AMP targets

A novel aspect of c-di-AMP signaling that also emerges from the present study is that *E. faecalis* reduces intracellular c-di-AMP pools when exposed to serum or urine (Fig. 3). Whether this is a relevant aspect of the adaptive process in the initial stages of infection or beyond remains to be determined. Salt concentrations in healthy human urine are relatively high (average osmolarity between 300 and 900 mosM/kg H_2_O) and low c-di-AMP levels have previously been linked to bacterial salt tolerance (59) whereas high c-di-AMP is associated with salt hypersensitivity (60, 61). It should be also noted that urine composition and osmolarity are constantly subjected to large fluctuations due to cycles of urine concentration and dilution, such that the ability of uropathogens to maintain osmotic balance in the bladder environment is (likely) an important aspect of bacterial UTI. On the other hand, serum osmolarity does not undego large fluctuations and normal osmolarity, ranging between 275 to 295 mosM/kg H_2_O, is considerably lower than urine. Thus, the environmental cues leading to the observed low levels of c-di-AMP in *E. faecalis* cells grown in urine or serum and the significance of this observation are unclear and awaits further investigation, particularly using *in vivo* infection models. Another possibility is that specific environmental cues present in serum and urine trigger c-di-AMP secretion, which could further deplete intracellular c-di-AMP. In *L. monocytogenes*, two multidrug efflux pumps were shown to export c-di-AMP from the cytosol during macrophage infection (62) such that a precedent of host-pathogen interactions triggering c-di-AMP secretion seems to already exist.

In bacteria featuring the DhhP/GdpP pair such as Streptococci and Staphylococci, the cytosolic DhhP is thought to play a dominant role in c-di-AMP degradation (10). While both DhhP and GdpP degrade c-di-AMP, DhhP cleaves c-di-AMP into the pApA intermediate and then further into the final hydrolysis product AMP (16, 18). GdpP, on the other hand, cleaves c-di-AMP into pApA only. Because DhhP converts pApA into AMP, it has been proposed that this activity is involved in feedback inhibition of GdpP-dependent c-di-AMP hydrolysis (63). Other than the cumulative effect of DhhP and GdpP in controlling intracellular c-di-AMP pools, loss of GdpP alone, for the most part, did not yield important phenotypes. In fact, the daptomycin MIC of Δ*gdpP* was identical to the parent strain, which is in an apparent contradiction with a report associating the emergence of a daptomycin-resistant *E. faecalis* isolate to a GdpP^I440S^ SNP that led to intracellular accumulation of c-di-AMP due to impaired GdpP activity (64). However, the GdpP^I440S^ strain harbored an additional point mutation in *liaR*, the response regulator of the LiaFSR system which is known for its critical regulatory role in cell membrane remodeling that is also important for daptomycin tolerance (65).

In the literature, there are several examples whereby c-di-AMP signaling modulates biofilm development and maturation, albeit the direction of the association of c-di-AMP with biofilm formation differs among bacterial species. For example, high levels of c-di-AMP due to inactivation of PDE-encoding genes led to reduced biofilms in *B. subtilis* and *Streptococcus gallolyticus* (66, 67) while to more robust biofilms in *S. mutans* and *S. pyogenes* (10, 47). Here, we showed that the Δ*cdaA* strain formed poor biofilms under *in vitro* or *in vivo* conditions (Fig. 5), which is in line with a recent report showing that a novel DAC inhibitor impairs biofilm formation and exopolysaccharide synthesis of *E. faecalis* (68). Here, we also found that expression of *ebpABC*, coding for the biofilm-associated and major virulence factor Ebp pilus, was strongly downregulated in the absence of c-di-AMP. Subsequent analyses revealed that, in addition to reduced mRNA/protein production, Ebp activity was severely compromised in the absence of c-di-AMP (Fig. 5). These results indicate that the adhesion/biofilm defect and attenuated virulence of the Δ*cdaA* strain can be also mattributed to defects in Ebp production, subunit translocation and pilus assembly. At this time, we propose that perturbations in cell envelope homeostasis hinders EbpABC biogenesis, surface anchoring or both, which would then explain why EbpA was undetected in the ELISA and the almost complete absence of pilus-like structures in the Δ*cdaA* strain (Fig. 5). It remains to be discovered how c-di-AMP regulates *ebpABC* transcription but initial studies should focus on probing possible interactions of c-di-AMP with known *ebpABC* transcriptional regulators such as EbpR, AhrC (also annotated as ArgR3), ArgR2, the Fsr quorum sensing system, and the RNA processing enzyme RNase J2 (69-72).

To evaluate the impact of c-di-AMP dysregulation at a global level, we used RNaseq to compare the transcriptome of *E. faecalis* OG1RF, Δ*dhhP*Δ*gdpP* and Δ*cdaA* strains. To our knowledge, this is the first time the transcriptome of a bona fide c-di-AMP^0^ strain was obtained, as previous global transcriptional analysis were restricted to strains with high c-di-AMP levels (9, 35, 67). In *B. subtilis*, c-di-AMP regulates transcription by interacting with a c-di-AMP-specific riboswitch (2) with several c-di-AMP riboswitch-regulated genes involved in K^+^ and osmolyte transport, and cell wall metabolism (73). While enterococcal genomes do not possess sequences that resemble the c-di-AMP riboswitch (2), several genes associated with osmolyte and oligopeptide transport were dysregulated (down or upregulated) in the Δ*cdaA* strain. At this time, it is unclear how c-di-AMP affects transcription of *E. faecalis* genes. One possibility is that an addidional (yet-to-be-identified) c-di-AMP riboswitch exist, as in the case of the structurally-related c-di-GMP which has been shown to bind two distinct c-di-GMP-specific riboswitches (74). A more plausible possibility that has been shown in other bacteria (7, 75-77), but does not invalidate the multiple riboswitch possibility, is that c-di-AMP allosterically controls the activity of transcriptional regulators that control transcription of transport-associated genes. Another important observation that emerges from the transcriptional analysis was the strong downregulation of genes from the ADI system on both Δ*cdaA* and Δ*pde*Δ*gdpP* transcriptomes. While the implications of this observation are unclear, it has been proposed that increases in arginine catabolism protects *E. faecalis* from oxidative stress and is linked to increased antibiotic tolerance (78). Also, the ADI operon is under control of two regulators of the ArgR family of transcriptional factors. Based on evidence that *ebpABC* transcription is regulated by two ArgR-type regulators (AhrC/Arg3 and Arg2) (70), future studies to explore a connection between c-di-AMP, arginine catabolism and Ebp expression are warranted.

In this report, we have added *E. faecalis* to the growing list of bacterial pathogens in which fitness, antibiotic tolerance and virulence is under c-di-AMP control. Most notably, we performed a systematic and unambiguous characterization of a genetically-stable *E. faecalis* c-di-AMP^0^ strain, which has not been conducted with any other bacterial species. Using the *Galleria mellonella* invertebrate model and two foreign body-associated mouse models, we showed that c-di-AMP is critical for the establishment of *E. faecalis* infections as the ability of strains that accumulated c-di-AMP to colonize and multiply in the host is severely compromised whereas the c-di-AMP^0^ strain is nonpathogenic. We also discovered that expression and biogenesis of the Ebp pilus was markedly impaired in the c-di-AMP^0^ strain, linking c-di-AMP signaling with expression of a major enterococcal virulence factor. Collectively, our results provide compelling evidence that approaches to interfere with c-di-AMP signaling might be highly effective for the treatment of *E. faecalis* infections, a pathogen of great medical concern due to its clinical prevalence and limited therapeutic options.

## MATERIALS AND METHODS

### Bacterial strains and growth conditions

The bacterial strains and plasmids used in this study are listed in Table 1. Strains were grown in brain heart infusion broth (BHI) or in chemically defined medium (CDM) originally developed for the growth oral streptococci (41) supplemented with 20 mM glucose at 37**°**C under static conditions. Strains harboring the pheromone-inducible pCIE plasmid (79) were grown under antibiotic pressure (10 μg ml^-1^ chloramphenicol) with or without the addition of 5 ng ml^-1^ ccF10 peptide (Mimotopes, USA). For growth kinetics assays, overnight cultures grown in CDM were adjusted to an OD_600_ of 0.25 and inoculated into fresh CDM or BHI at a ratio of 1:50. Growth was monitored at OD_600_ using an automated growth reader (Oy Growth Curves AB Ltd, Finland). NaCl, KCl, peptone, glycine betaine hydrochloride or carnitine (all purchased from Sigma Aldrich, USA) and c-di-AMP (InvivoGen, USA) were added to the growth media at the concentrations indicated in the figure legends.

### General cloning techniques

The nucleotide sequences of *cdaA, dhhP* and *gdpP* were obtained from the *E. faecalis* OG1RF genome via BioCyc (80). The Wizard genomic DNA purification kit (Promega, Madison, WI) was used for isolation of bacterial genomic DNA (gDNA), and the Monarch plasmid miniprep kit (New England BioLabs, Ipswich, MA) used for plasmid purification. The Monarch DNA gel extraction kit (New England BioLabs) was used to isolate PCR products. Colony PCR was performed using PCR 2x Master Mix (Promega) with primers listed in Table S3.

### Construction of gene deletions and genetically-complemented strains

**Deletion** of *cdaA*, *dhhP*, and *gdpP* from *E*. *faecalis* OG1RF was carried out using the pCJK47 markerless genetic exchange system (40) as previously described. The upstream and downstream sequences of *cdaA*, *dhhP*, and *gdpP* were amplified using the primers listed in Table S3. Introduction of amplicons into the pCJK47 vector, followed by conjugation into *E*. *faecalis* OG1RF and isolation of deletion mutants were carried out as described elsewhere (40), but using CDM to grow the Δ*cdaA* strain. The Δ*dhhP*Δ*gdpP* double mutant was obtained by conjugating the pCJK-*gdpP* plasmid with the Δ*dhhP* mutant. All gene deletions were confirmed by PCR sequencing of the insertion site and flanking sequences. The full-length *cdaA* gene was amplified by PCR using the primers listed in Table S3, digested with the appropriate restriction enzymes, and ligated into pCIE vector to yield plasmid pCIE::*cdaA*. The plasmid was propagated in *E. coli* DH10b, verified by sequencing and electroporated into the *E. faecalis* Δ*cdaA* strain as described elsewhere (81).

### Antibiotic susceptibility assay

Strains were grown overnight in CDM, normalized to an OD_600_ of 0.5, and diluted 1:1000. The diluted cultures were inoculated into fresh medium containing increasing concentrations of antibiotics (ampicillin, bacitracin, daptomycin, or vancomycin; all purchased from Sigma Aldrich) at a ratio of 1:20, and incubated at 37 °C for 24 hrs. The absorbance at OD_600_ was measured in a Synergy H1 microplate reader (Molecular Devices, USA).

### *Ex vivo* survival in serum and urine

Strains were grown overnight in CDM, normalized to an OD_600_ of 0.5, and inoculated at a 1:1000 ratio into pooled human serum or pooled human urine (both purchased from Lee Biosolutions, USA). At selected time points, aliquots were serially diluted in PBS and the dilutions plated on CDM agar for CFU determination.

### c-di-AMP quantifications

Overnight cultures grown in CDM were diluted 1:20 in fresh CDM and grown until the cultures reached an OD_600_ of ∼ 0.5. The cultures were immediately filtered through a 0.45-micron filter (Millipore, USA), and the cells collected on the filter membrane transferred into a tube containing ice-cold PBS using a cell scraper (Biologix, USA). Cell pellets were obtained by centrifugation at 4000 g for 5 mins and resuspended into cold extraction buffer (acetonitrile/MeOH/H_2_0 – 50:40:10; LC-MS grade) adding purified c-di-GMP (Sigma Aldrich) as an internal standard. Cell suspensions were frozen with liquid nitrogen for 15 min, and then incubated at 95°C for 10 mins. Samples were mixed with 0.5 ml of 1 mm glass beads and lysed in a bead beater thrice at 45 secs intervals. Glass beads and cell debris were pelleted by centrifugation at 20,000 g for 20 mins at 4 °C, and the supernatants stored overnight at -20°C. The next day, the nucleotide extracts were centrifuged at 10,000 g for 30 mins to remove protein precipitate, and the samples dried at 40°C using nitrogen gas and then stored at 4°C. The dried samples were resuspended into 300 μl extraction buffer and passed through solid-phase extraction (SPE) column. The filtrates were analyzed by LC-MS/MS (LCMS-8060; triple quadrupole system) at PhenoSwitch Bioscience, Inc (Quebec, Canada). The values obtained were normalized by total milligram of protein that was determined using the Bradford assay.

### Biofilm assay

Overnight cultures grown in CDM were normalized to OD_600_ of 0.5, and further diluted 1:25 in fresh CDM supplemented with 0.175% glucose and added to the wells of 96-well polystyrene plates (Grenier CellSTAR, USA) that were then incubated at 37°C under static conditions for 24 hrs. After incubation, media containing planktonic cells was discarded from the wells, which were gently washed twice with PBS. Adherent cells were stained with 0.1% crystal violet for 15 min, and the bound dye eluted in a 33% acetic acid solution. Absorbance was measured at an OD_595_ using the Synergy H1 microplate reader (Biotek, USA).

### Adhesion assay

Pili-mediated adhesion assay was performed as previously described (82) with minor modifications. Briefly, 96-well polystyrene plates (Grenier CellSTAR) either uncoated or coated with 200 μL of bovine serum albumin (BSA) (200 μg ml^-1^) overnight at 4°C were used. Coated microtiter plates were washed twice with 200 μl PBS and blot dried. Cultures were grown overnight at 37 °C in 10 ml CDM, washed with PBS and normalized to OD_600_ of 0.5. Then, 100 μl of normalized bacterial culture were added to wells of the microtiter plate, and 50 mM Na_2_CO_3_ suspended in PBS was added to induce pilus expression as previously described (52). The cells were incubated at 37°C for 2 hrs, washed twice with 200 μl PBS and stained with 0.1% (w/v) crystal violet as described previously. Absorbance was measured at an optical density at 595 nm using the Synergy H1 microplate reader.

### G. mellonella infection

Larvae of *G*. *mellonella* were used as model to assess virulence of *E. faecalis* strains as described previously (34) with minor modifications. Briefly, groups of 15 larvae (200–300 mg in weight) were injected with 5 μl of bacterial inoculum containing ∼1x10^5^ CFU. Larvae injected with heat-inactivated *E*. *faecalis* (30 mins at 100°C) or PBS were used as negative and vehicle controls, respectively. Post-injection, larvae were kept at 37°C and survival recorded at selected intervals for up to 96 hours.

### Mouse CAUTI model

The methods for the CAUTI experiments have been published (82). Thus, only a general overview and minor modifications are presented here. Female C57BL/6Ncr 6-weeks old mice (Charles River Laboratories, USA) were anesthetized and subjected to transurethral implantation of a 5-mm length platinum-cured silicone catheter. Immediately after catheter implantation, mice were infected with **∼**1x10^7^ CFU of bacteria in PBS. 24 hrs post-infection, mice were euthanized and the catheter and organs collected. CFU enumeration was performed by plating catheter and organ homogenates on CDM agar plates selective for *E. faecalis* OG1RF and derivatives (10 μg ml^-1^ fusidic acid and 200 μg ml^-1^ rifampicin). This procedure was approved and performed in compliance with the University of Notre Dame Institutional Animal Care and Use Committee (IACUC).

### Foreign body-associated peritonitis mouse model

The methods for the foreign body-associated peritonitis model have also been published (49), such that only a general overview and minor modifications are described here. The day before surgey, 1-cm long segments of a silicone catheter tubing (Qosina, USA) were cut and coated in 100 µg ml**^-1^** human fibrinogen solution (Sigmna Aldrich) at 4°C overnight. The following day, female C57BL/6J 8-week old mice (Jackson Laboratories, USA) were anesthetized and one 1-cm long catheter segment inserted into the peritoneum using an 18-gauge BD Spinal needle/stylette (Becton, Dickinson and Company, USA). Four hrs post-catheter implantation, mice were injected via i.p. with ∼2×10^8^ CFU of bacteria in PBS. Animals were euthanized 48 hrs post-infection and peritoneal wash, catheter and spleen collected for CFU enumeration by plating serial dilutions on CDM agar plates selective for *E. faecalis* OG1RF and derivatives. This procedure was approved and performed in compliance with the University of Florida Institutional Animal Care and Use Committee (IACUC).

### Electron microscopy analyzes

Transmission electron microscopy (TEM) and scanning electron microscopy (SEM) were performed at the Interdisciplinary Center for Biotechnology Research (ICBR) Electron Microscopy core lab of the University of Florida. SEM and TEM analyses were performed using standard preparation and visualization procedures. For immunogold labelled TEM analysis, nickel grids (400 Ni-UB) were coated with poly-L-lysine solution at a dilution of 1:10 for 15 mins and air-dried. Stationary phase *E. faecalis* was diluted at 1:1000 and 5 µl of the diluent was pipetted on the surface of the grid and incubated for 5 mins. The bacterial cells were fixed with 4% (w/v) paraformaldehyde for 20 min, washed with PBS and blocked with skimmed milk for 30 min. After blocking, the grids were washed with PBS thrice, and incubated with primary mouse anti-EbpA^Full^ antibody (82) diluted 1:1000 for 1 hr at room temperature. Post-incubation, the grids were washed with PBS thrice and incubated with 12-nm gold donkey anti-goat antibody (1:20 dilution) for 1 hr at room temperature. The grids were washed with PBS thrice, and then washed with sterile water before being negatively stained with 0.02% uranyle acetate for 10 secs, and air-dried before imaging.

### ELISA and immunoblot analyses

Surface expression of EbpA by *E. faecalis* OG1RF and mutant strains was determined by ELISA as previously described (83). Samples were prepared for immunoblotting (51, 53) with minor modifications. Briefly, cultures were grown overnight in CDM, diluted 1:20 into fresh CDM, and grown to an OD_600_ of 0.5. Cell pellets were collected by centrifugation, washed with PBS and incubated with 50 µl of a 30 mg ml**^-1^** lysozyme solution (Sigma Aldrich) for 20 mins to separate cell wall and protoplast fractions. Lysozyme-treated cells were centrifuged at 4,000 g for 10 mins at 4°C. The supernatant collected represents the cell wall fraction while the cell pellets collected represent the protoplast fraction. Protein extracts were resuspended in 2X SDS buffer with 2.5% beta-merceptenthanol and 0.1 M dithiothreitol (DTT) and then boiled at 100°C for 15 mins prior to loading onto 6% separating tris-glycine gel. Immunoblotting was performed using wet transfer to PVDF membranes in tris-glycine transfer buffer at 100 V for 90 mins at 4°C. After the transfer was complete, the PVDF membrane was soaked overnight in blocking buffer (PBS, 0.05% Tween-20, 3% BSA) at 4°C with constant agitation. The next day, the membrane was washed thrice with washing buffer (PBS, 0.05% Tween-20) followed by 1 hr incubations with anti-EbpA^FULL^ antibody (1:500) in blocking buffer, secondary goat anti-mouse antibody (Invitrogen, USA) (1:4000) with washes before and after each incubation. Proteins were visualized using the ECL detection kit (Amersham, USA) in a ChemiDoc imaging system (Bio-Rad, USA). As loading control, a separate blot was prepared in a similar manner using a rabbit polyclonal anti-*S. pyogenes* DnaK (1:1000) antibody (84).

### RNA analysis

Total RNA was isolated from *E. faecalis* cells grown to OD_600_ = 0.35 by acid-phenol-chloroform extractions and preparation of cDNA libraries were performed as previously described (85). Sample quality and quantity were assessed on an Agilent 2100 Bioanalyzer at the University of Florida ICBR. RNA deep sequencing (RNAseq) was performed at UF-ICBR using the Illumina NextSeq 500 platform. Read mapping was performed on a Galaxy server hosted by the University of Florida Research Computer using Map with Bowtie for Illumina and the *E. faecalis* OG1RF genome (GenBank accession no. CP002621.1) used as reference. The reads per open reading frame were tabulated with htseq-count. Final comparisons between parent and mutant strains were performed with Degust (http://degust.erc.monash.edu/), with a false-discovery rate (FDR) of 0.05 and a 2-fold change cutoff. Quantifications of *cdaA*, *dhhP*, *gdpP* and *ebpA* mRNA were obtained by quantitative real-time PCR (qRT-PCR) as previously described (85).

### Statistical analysis

Data were analysed using GraphPad Prism 8.0 software (GraphPad Software, San Diego, CA, USA). Data from multiple experiments were pooled, and appropriate statistical tests were applied, as indicated in the figure legends. An adjusted value of *p* ≤ 0.05 was considered statistically significant. * *p* ≤ 0.05, ** *p* ≤ 0.01, *** *p* ≤ 0.001, **** *p* ≤ 0.0001.

## Data availability

Gene expression data have been deposited in the NCBI Gene Expression Omnibus (GEO) database (https://www.ncbi.nlm.nih.gov/geo) under GEO Series accession number GSE174381.

## Acknowledgements

This study was supported by NIH-NIAID R21 AI135158 to J.A.L. and by by institutional funds from the University of Notre Dame to A.L.F.M. L.G.C was supported by NIH-NIDCR Training Grant T90 DE021990.

---

**FIG S1.**
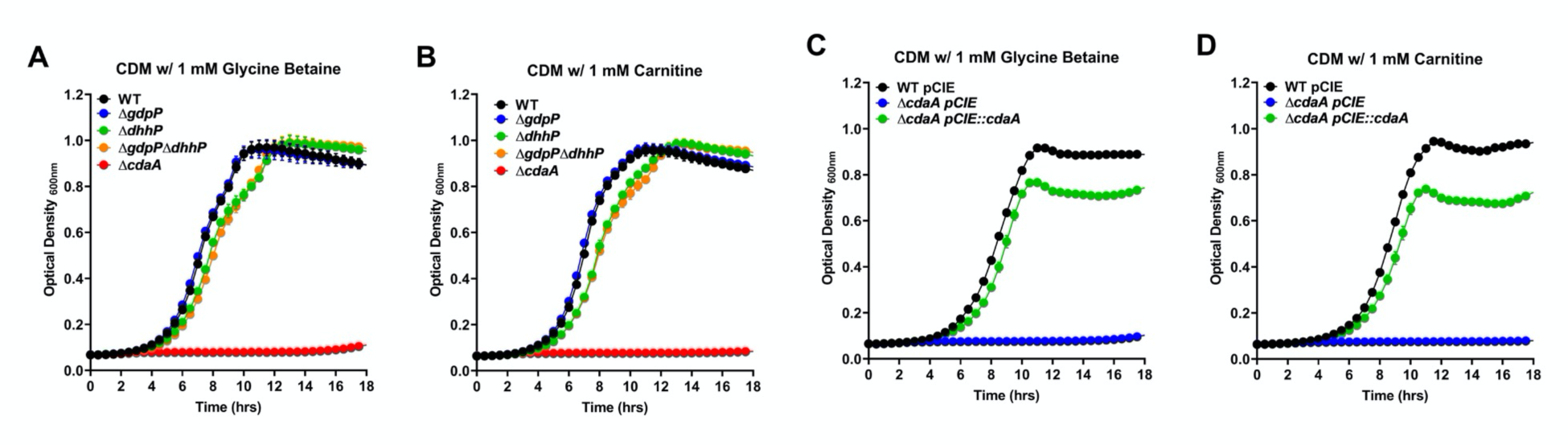
Growth of *E. faecalis* OG1RF (WT) and Δ*cdaA* strains in CDM containing increasing concentrations of NaCl or KCl. Growth curves in CDM supplemented with 0.01% peptone with increasing concentration of NaCl (A-B) or KCl (C-D). Growth curves in CDM with increasing concentration of NaCl (E) or KCl (F). Data points represent nine independent biological replicates and error bars represent the standard error of margin.

**FIG S2.**
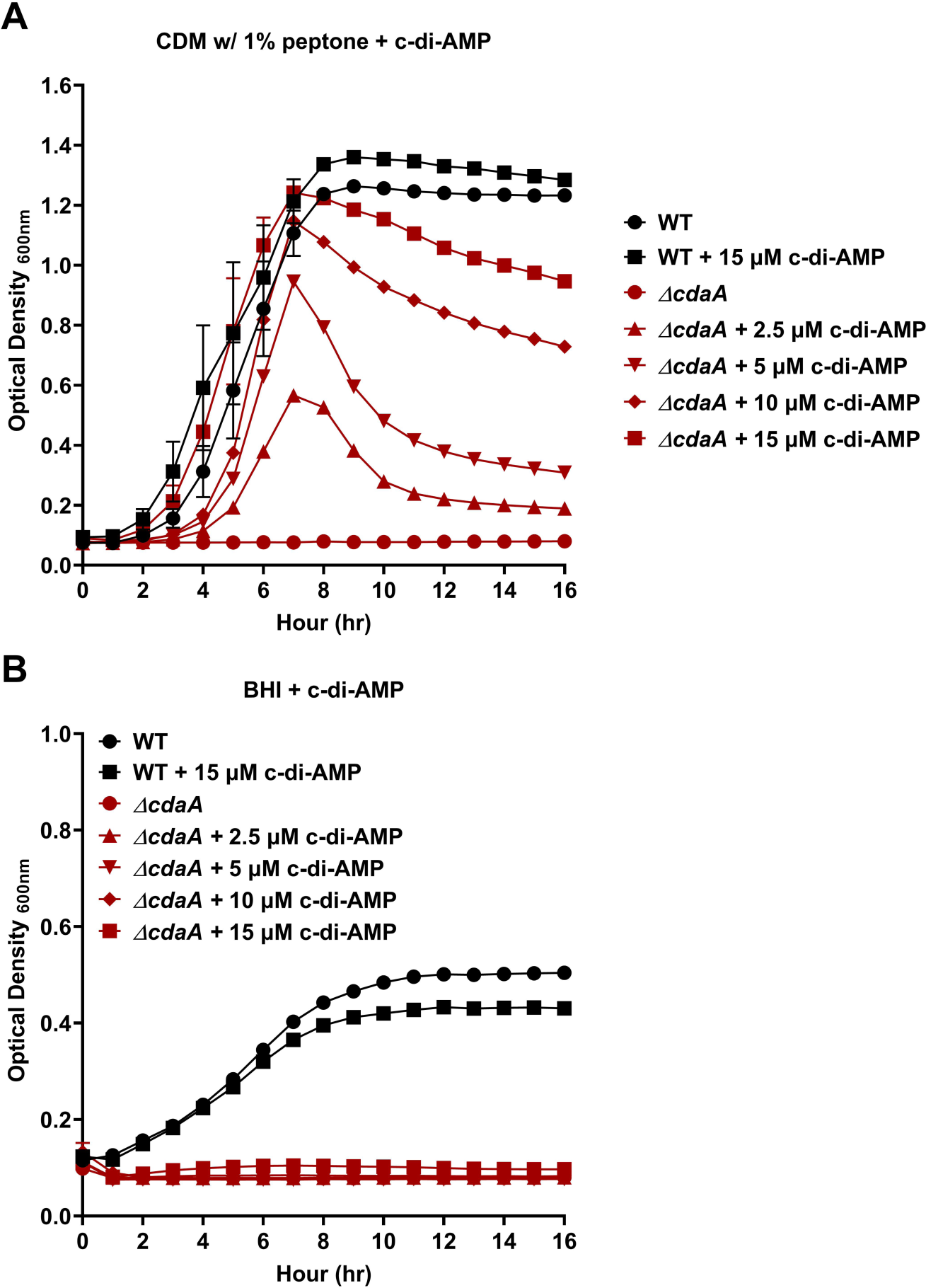
Growth of *E. faecalis* OG1RF (WT) and Δ*cdaA* strains in CDM + 0.1% peptone (A) or BHI (B) containing increasing concentrations of purified c-di-AMP.

**FIG S3.**
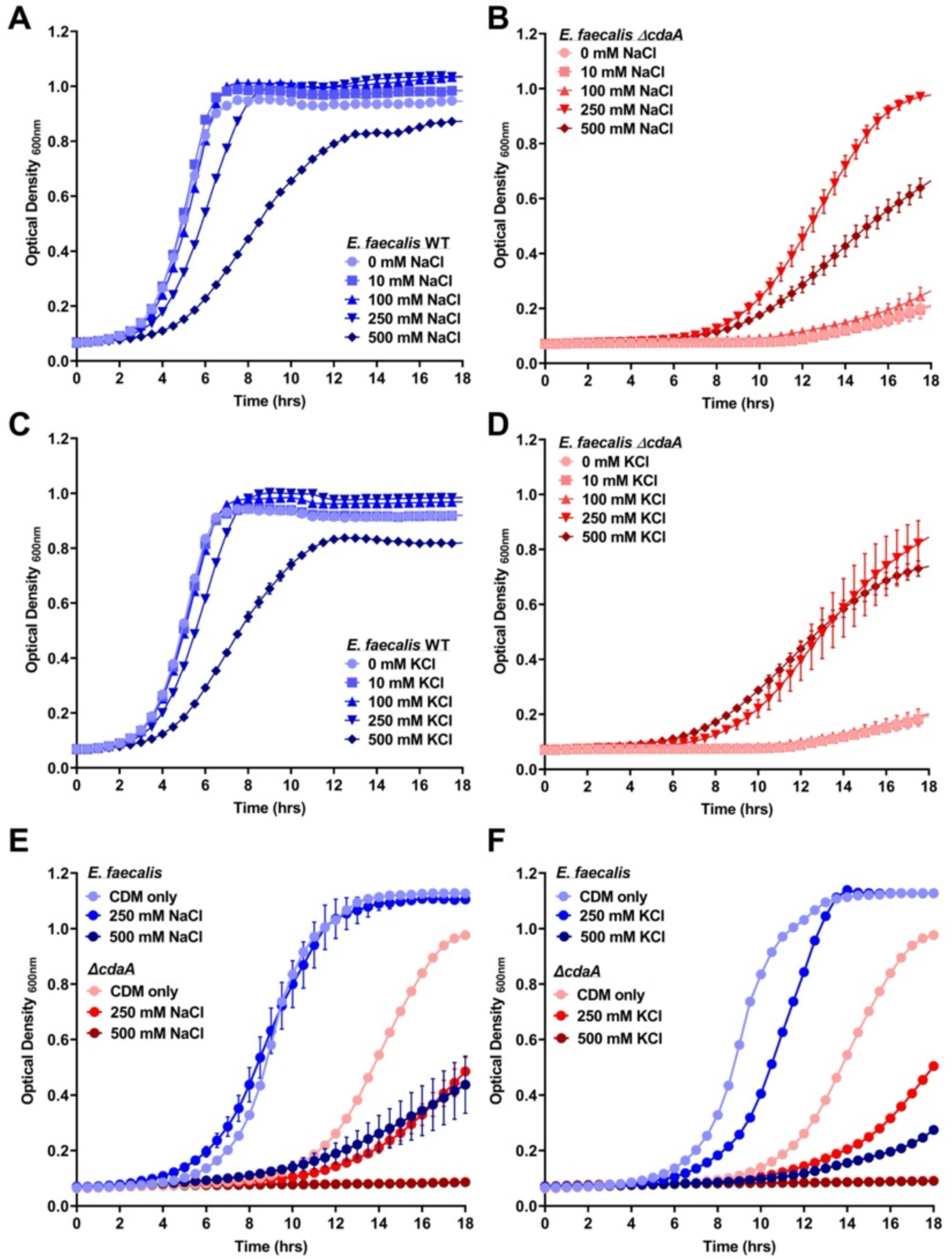
Growth of *E. faecalis* OG1RF (WT) and Δ*cdaA* strains in CDM containing increasing concentrations of NaCl or KCl. Growth curves in CDM supplemented with 0.01% peptone with increasing concentration of NaCl (A-B) or KCl (C-D). Growth curves in CDM with increasing concentration of NaCl (E) or KCl (F). Data points represent nine independent biological replicates and error bars represent the standard error of margin.

**FIG S4.**
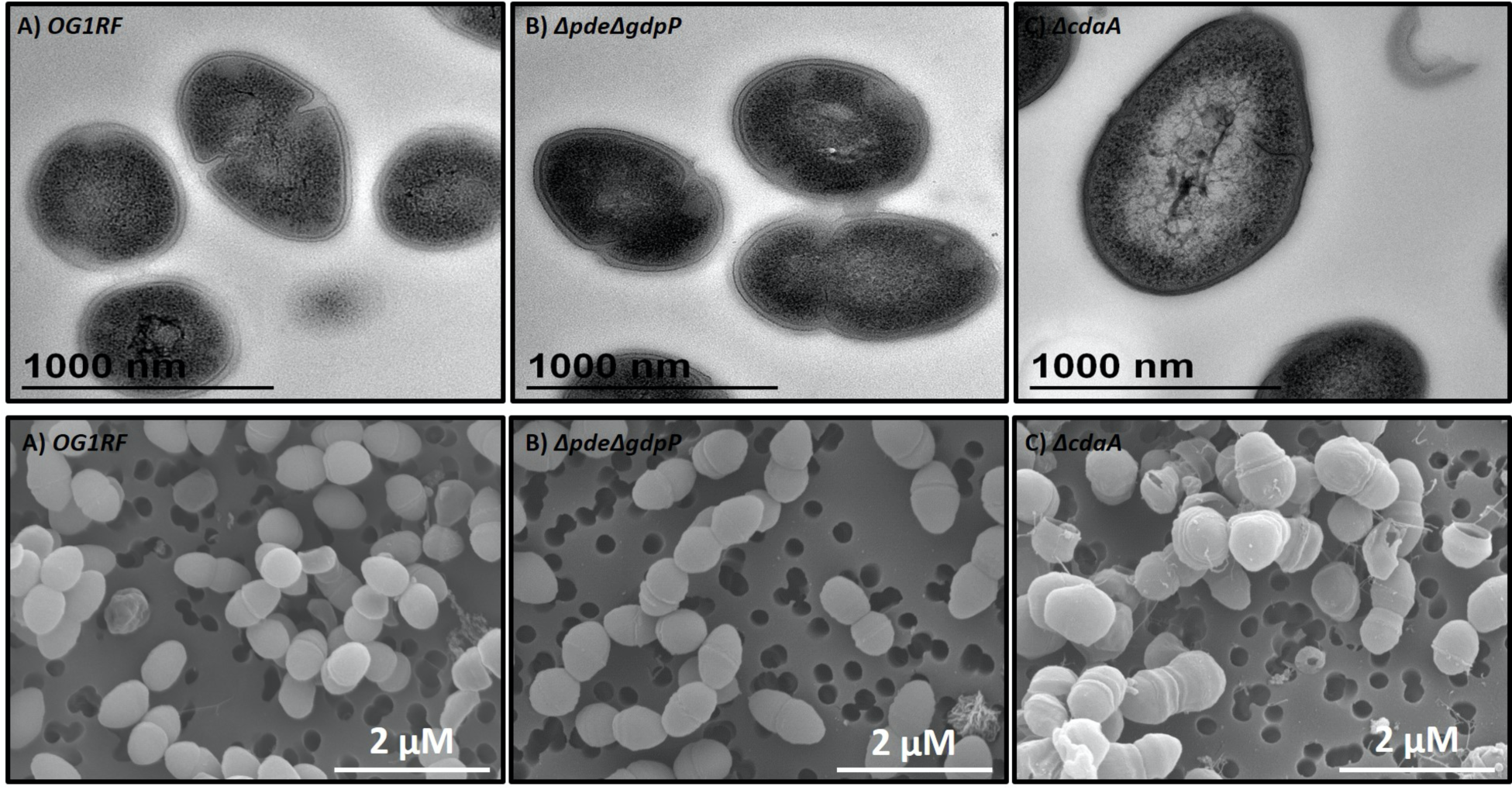
Electron microscopy images of *E. faecalis* OG1RF, Δ*pde*Δ*gdpP* and Δ*cdaA* strains. Top: Representative transmission electron microscopy (TEM) images. Bottom: Representative scanning electron microscope (SEM) images.

## Other Supporting information

**TABLE S1** List of genes differentially expressed (FDR of 0.05 and 2-fold change cutoff) in *E. faecalis* Δ*cdaA* strain in comparison to OG1RF (wild type) strain.

**TABLE S2** List of genes differentially expressed (FDR of 0.05 and 2-fold change cutoff) in *E. faecalis* Δ*dhhPΔgdpP* in comparison to OG1RF (wild type) strain.

**TABLE S3** Primers used in this study.

