## Supplemental Table 1 for "c-di-AMP is essential for the virulence of *Enterococcus faecalis*"

**Table S1.** List of genes differentially expressed (FDR of 0.05 and 2-fold change cutoff) in *E. faecalis* ∆*cdaA* strain in comparison to OG1RF (wild type) strain.

| **Old Gene Locus** | **New Gene Locus** | **Log2FC** | **P-Value** | **Annotated Function** | **Functional Category** |
| --- | --- | --- | --- | --- | --- |
| OG1RF_10004 | OG1RF_RS00030 | 1.0191 | 7.14E-12 | DNA replication and repair protein RecF | DNA replication and repair |
| OG1RF_10017 | OG1RF_RS00090 | 1.4074 | 6.58E-15 | levR/ transcriptional regulatory protein LevR | DNA replication and repair |
| OG1RF_10030 | OG1RF_RS00155 | 3.0606 | 1.34E-09 | hypothetical protein | Function unknown |
| OG1RF_10032 | OG1RF_RS00165 | 1.1165 | 2.30E-03 | hypothetical protein | Function unknown |
| OG1RF_10057 | OG1RF_RS00290 | 3.4022 | 8.84E-20 | oligopeptide ABC superfamily ATP binding cassette transporter, binding protein | Peptide ABC transporter |
| OG1RF_10058 | OG1RF_RS00295 | 3.7649 | 2.73E-10 | hypothetical protein | Function unknown |
| OG1RF_10059 | OG1RF_RS00300 | 3.8326 | 1.98E-14 | LLM class flavin-dependent oxidoreductase | Respiration and ATP metabolism |
| OG1RF_10060 | OG1RF_RS00305 | 1.9126 | 4.77E-14 | ruvA/crossover junction ATP-dependent DNA helicase | DNA replication and repair |
| OG1RF_10061 | OG1RF_RS00310 | 1.1726 | 1.55E-12 | ruvB/crossover junction ATP-dependent DNA helicase | DNA replication and repair |
| OG1RF_10062 | OG1RF_RS00315 | 1.5269 | 1.04E-07 | hypothetical protein | Function unknown |
| OG1RF_10063 | OG1RF_RS00320 | 1.0423 | 2.45E-08 | N-acylglucosamine-6-phosphate 2-epimerase | Central metabolism |
| OG1RF_10068 | OG1RF_RS00345 | -1.1908 | 1.46E-12 | 3-oxoacyl-[acyl-carrier-protein] reductase | Central metabolism |
| OG1RF_10077 | OG1RF_RS00390 | 1.3919 | 1.10E-06 | atpF/ ATP sunthase F0 sector subunit B | Respiration and ATP metabolism |
| OG1RF_10078 | OG1RF_RS00395 | 1.8302 | 1.10E-06 | hypothetical protein | Function unknown |
| OG1RF_10089 | OG1RF_RS00445 | -1.4798 | 2.57E-12 | formate/nitrite transporter | Central metabolism |
| OG1RF_10097 | OG1RF_RS00485 | -2.1706 | 7.33E-11 | ArgR family transcriptional regulator | Central metabolism |
| OG1RF_10098 | OG1RF_RS00490 | -4.9226 | 3.00E-16 | argR/arginine repressor | Central metabolism |
| OG1RF_10099 | OG1RF_RS00495 | -5.1278 | 9.65E-17 | arcA/arginine deiminase | Central metabolism |
| OG1RF_10100 | OG1RF_RS00500 | -5.5563 | 1.23E-15 | arcB/ornithine carbamoyltransferase | Central metabolism |
| OG1RF_10101 | OG1RF_RS00505 | -4.1165 | 7.22E-15 | arcC/carbamate kinase | Central metabolism |
| OG1RF_10102 | OG1RF_RS00510 | -3.5767 | 1.05E-14 | ntcA/global nitrogen regulator NtcA | Central metabolism |
| OG1RF_10103 | OG1RF_RS00515 | -3.5406 | 2.13E-17 | C4-dicarboxylate ABC transporter | Peptide ABC transporter |
| OG1RF_10155 | OG1RF_RS00780 | -1.0385 | 6.89E-09 | rpsS/30S ribosomal protein S19 | DNA replication and repair |
| OG1RF_10181 | OG1RF_RS00910 | 1.2201 | 9.18E-12 | M20D family peptidase | Central metabolism |
| OG1RF_10188 | OG1RF_RS00945 | 2.3224 | 4.13E-12 | GNAT family acetyltransferase | Central metabolism |
| OG1RF_10191 | OG1RF_RS00960 | -1.1155 | 1.89E-13 | amino acid ABC superfamily ATP binding cassette transporter, membrane protein | Peptide ABC transporter |
| OG1RF_10198 | OG1RF_RS00990 | 1.0556 | 3.30E-06 | aldA/aldehyde dehydrogenase (NAD(+)) | Respiration and ATP metabolism |
| OG1RF_10216 | OG1RF_RS01220 | -2.5795 | 2.10E-04 | bglA/6-phospho-beta-glucosidase | Central metabolism |
| OG1RF_10217 | OG1RF_RS01225 | -2.2013 | 2.68E-03 | phosphoglycerate mutase | Central metabolism |
| OG1RF_10246 | OG1RF_RS01375 | 1.2925 | 1.09E-08 | flavin reductase domain protein, FMN-binding protein | Respiration and ATP metabolism |
| OG1RF_10255 | OG1RF_RS01420 | 1.4333 | 4.82E-15 | lysC/ aspartate kinase | Central metabolism |
| OG1RF_10272 | OG1RF_RS01505 | 1.2817 | 2.25E-07 | hypothetical protein | Respiration and ATP metabolism |
| OG1RF_10273 | OG1RF_RS01510 | 1.3464 | 2.02E-08 | major facilitator family transporter | Multifunction transporter |
| OG1RF_10274 | OG1RF_RS01515 | 1.1654 | 4.74E-07 | arcC2/carbamate kinase 2 | Central metabolism |
| OG1RF_10304 | OG1RF_RS01670 | 1.7216 | 3.61E-07 | adenylosuccinate synthetase | Central metabolism |
| OG1RF_10305 | OG1RF_RS01675 | 1.1478 | 1.42E-07 | hypothetical protein | Function unknown |
| OG1RF_10310 | OG1RF_RS01700 | 1.4458 | 7.56E-07 | eda-1/2-dehydro-3-deoxy-phosphogluconate aldolase | Central metabolism |
| OG1RF_10311 | OG1RF_RS01705 | 1.6618 | 4.25E-10 | PfkB family carbohydrate kinase | Central metabolism |
| OG1RF_10312 | OG1RF_RS01710 | 1.7366 | 7.39E-11 | kduI1/4-deoxy-L-threo-5-hexosulose-uronate ketol-isomerase | Central metabolism |
| OG1RF_10313 | OG1RF_RS01715 | 1.9769 | 3.21E-11 | idnO/gluconate 5-dehydrogenase | Respiration and ATP metabolism |
| OG1RF_10326 | OG1RF_RS01780 | -5.2800 | 8.88E-17 | dtPT/POT family proton (H+)-dependent oligopeptide transporter | Peptide ABC transporter |
| OG1RF_10328 | OG1RF_RS01790 | -1.6351 | 1.96E-12 | peptidase M23 | Central metabolism |
| OG1RF_10337 | OG1RF_RS01835 | -1.0464 | 5.93E-12 | acetate CoA ligase | Respiration and ATP metabolism |
| OG1RF_10354 | OG1RF_RS01920 | 1.3665 | 1.41E-11 | hypothetical protein | Function unknown |
| OG1RF_10357 | OG1RF_RS01935 | -1.1826 | 9.31E-10 | class Ib ribonucleoside-diphosphate reductase assembly flavoprotein NrdI | Respiration and ATP metabolism |
| OG1RF_10358 | OG1RF_RS01940 | -1.6191 | 2.75E-09 | nrdH/glutaredoxin | Respiration and ATP metabolism |
| OG1RF_10377 | OG1RF_RS02040 | 1.3755 | 2.00E-07 | C4-dicarboxylate ABC transporter | Peptide ABC transporter |
| OG1RF_10378 | OG1RF_RS02045 | 1.2930 | 7.22E-08 | galE/NAD-dependent epimerase/dehydratase | Central metabolism |
| OG1RF_10379 | OG1RF_RS02050 | 1.0913 | 1.30E-05 | phage integrase family site-specific recombinase | DNA replication and repair |
| OG1RF_10380 | OG1RF_RS02055 | 1.2935 | 9.40E-07 | nitroreductase | Central metabolism |
| OG1RF_10386 | OG1RF_RS02085 | 1.2102 | 2.00E-09 | nitroreductase | Central metabolism |
| OG1RF_10387 | OG1RF_RS02090 | 1.6670 | 1.23E-11 | glyoxalase | Central metabolism |
| OG1RF_10395 | OG1RF_RS02130 | 1.1042 | 1.60E-05 | hypothetical protein | Function unknown |
| OG1RF_10425 | OG1RF_RS02270 | -0.9750 | 9.45E-07 | histidine triad protein | Respiration and ATP metabolism |
| OG1RF_10446 | OG1RF_RS02375 | 1.0625 | 5.52E-07 | hypothetical protein | Function unknown |
| OG1RF_10473 | OG1RF_RS02510 | -0.9862 | 3.24E-10 | amidase | Central metabolism |
| OG1RF_10474 | OG1RF_RS02515 | -1.7635 | 5.10E-12 | NMN family nicotinamide monnucleotide uptake permease | Pyridine and purine metabolism |
| OG1RF_10475 | OG1RF_RS02525 | -1.2122 | 1.19E-09 | deoxynucleoside kinase | Respiration and ATP metabolism |
| OG1RF_10481 | OG1RF_RS02555 | 1.9771 | 4.02E-14 | serine hydrolase | Central metabolism |
| OG1RF_10482 | OG1RF_RS02560 | 1.5108 | 1.45E-11 | hypothetical protein | Function unknown |
| OG1RF_10485 | OG1RF_RS02570 | -1.4173 | 1.64E-10 | cell wall surface anchor protein | Cell surface adhesin |
| OG1RF_10486 | OG1RF_RS02575 | -1.7211 | 8.30E-12 | WxL domain surface protein | Cell surface adhesin |
| OG1RF_10487 | OG1RF_RS02580 | -1.5892 | 4.15E-11 | WxL domain surface protein | Cell surface adhesin |
| OG1RF_10488 | OG1RF_RS02585 | -1.5886 | 5.50E-14 | legume lectins beta domain protein | Central metabolism |
| OG1RF_10489 | OG1RF_RS02590 | -1.5385 | 3.02E-10 | WxL domain surface protein | Cell surface adhesin |
| OG1RF_10490 | OG1RF_RS02595 | -1.0796 | 1.92E-09 | cell wall surface anchor family protein | Cell surface adhesin |
| OG1RF_10491 | OG1RF_RS02600 | -1.3591 | 1.33E-09 | hypothetical protein | Function unknown |
| OG1RF_10499 | OG1RF_RS02640 | 1.5523 | 2.10E-08 | hypothetical protein | Function unknown |
| OG1RF_10516 | OG1RF_RS02725 | 1.0861 | 7.50E-13 | acyltransferase | Central metabolism |
| OG1RF_10536 | OG1RF_RS02820 | -1.4936 | 2.42E-08 | hypothetical protein | Function unknown |
| OG1RF_10537 | OG1RF_RS02825 | -1.3523 | 1.70E-06 | aatB/amino acid ABC superfamily ATP binding cassette transporter, binding protein | Peptide ABC transporter |
| OG1RF_10538 | OG1RF_RS02830 | -1.6843 | 2.77E-08 | ABC superfamily ATP binding cassette transporter, ABC protein | Peptide ABC transporter |
| OG1RF_10539 | OG1RF_RS02835 | -1.6837 | 1.70E-06 | amino acid ABC superfamily ATP binding cassette transporter, membrane protein | Peptide ABC transporter |
| OG1RF_10551 | OG1RF_RS02890 | 1.6848 | 1.47E-11 | GTP cyclohydrolase | Respiration and ATP metabolism |
| OG1RF_10552 | OG1RF_RS02895 | 1.3718 | 8.23E-12 | rplY/50S ribosomal protein L25 | Protein synthesis, translation and degradation |
| OG1RF_10556 | OG1RF_RS02920 | -1.0999 | 2.90E-11 | udk/uridine kinase | Pyridine and purine metabolism |
| OG1RF_10557 | OG1RF_RS02925 | 1.4753 | 1.30E-05 | mRNA interferase PemK | DNA replication and repair |
| OG1RF_10580 | OG1RF_RS03040 | 1.0870 | 1.83E-10 | mRNA interferase PemK | DNA replication and repair |
| OG1RF_10591 | OG1RF_RS03095 | 1.1228 | 3.74E-09 | N-acetyltransferase | Central metabolism |
| OG1RF_10597 | OG1RF_RS03120 | 1.4251 | 5.36E-10 | glyoxalase | Central metabolism |
| OG1RF_10600 | OG1RF_RS03135 | 1.8883 | 3.55E-16 | calcium-translocating P-type ATPase, PMCA-type | Multifunction transporter |
| OG1RF_10601 | OG1RF_RS03140 | -1.0861 | 5.40E-13 | potassium transporter Kup | Multifunction transporter |
| OG1RF_10627 | OG1RF_RS03265 | -0.9864 | 8.95E-11 | bifunctional acetaldehyde-CoA/alcohol dehydrogenase | Respiration and ATP metabolism |
| OG1RF_10655 | OG1RF_RS03405 | -2.1110 | 4.94E-14 | glucose transporter GlcU | Central metabolism |
| OG1RF_10656 | OG1RF_RS03410 | -1.2168 | 1.12E-11 | amino acid permease | Central metabolism |
| OG1RF_10667 | OG1RF_RS03465 | -1.4375 | 3.82E-13 | folate ECF transporter | Multifunction transporter |
| OG1RF_10671 | OG1RF_RS03485 | -1.6636 | 9.29E-15 | extracellular protein | Cell surface adhesin |
| OG1RF_10679 | OG1RF_RS03525 | 1.4942 | 7.13E-08 | hypothetical protein | Function unknown |
| OG1RF_10700 | OG1RF_RS03625 | 1.3272 | 1.40E-05 | hypothetical protein | Function unknown |
| OG1RF_10707 | OG1RF_RS03660 | 2.7053 | 3.24E-14 | DNA repair exonuclease | DNA replication and repair |
| OG1RF_10721 | OG1RF_RS03735 | 1.1405 | 4.11E-12 | division/cell wall cluster transcriptional repressor MraZ | Central metabolism |
| OG1RF_10722 | OG1RF_RS03740 | 1.0532 | 3.91E-12 | 16S rRNA (cytosine(1402)-N(4))-methyltransferase | Protein synthesis, translation and degradation |
| OG1RF_10723 | OG1RF_RS03745 | 1.2195 | 3.08E-11 | cell division protein FtsL | Central metabolism |
| OG1RF_10739 | OG1RF_RS03825 | 1.2569 | 4.60E-05 | DUF4828 domain-containing protein | Function unknown |
| OG1RF_10745 | OG1RF_RS03850 | -2.2074 | 1.20E-07 | PTS sugar transporter subunit IIB | Central metabolism |
| OG1RF_10750 | OG1RF_RS03875 | -2.9269 | 7.44E-10 | PTS sugar transporter subunit IIB | Central metabolism |
| OG1RF_10751 | OG1RF_RS03880 | -2.0275 | 2.31E-08 | PTS mannose transporter subunit IIA | Central metabolism |
| OG1RF_10770 | OG1RF_RS03980 | 1.1822 | 4.01E-13 | aspartate 4-decarboxylase | Central metabolism |
| OG1RF_10771 | OG1RF_RS03985 | 1.2709 | 4.67E-09 | putative lipoprotein | Cell surface adhesin |
| OG1RF_10796 | OG1RF_RS04190 | -1.2734 | 5.56E-07 | hypothetical protein | Function unknown |
| OG1RF_10797 | OG1RF_RS04195 | -1.0786 | 2.58E-07 | recN2/DNA repair protein RecN | DNA replication and repair |
| OG1RF_10798 | OG1RF_RS04200 | -1.0136 | 4.31E-03 | hypothetical protein | Function unknown |
| OG1RF_10805 | OG1RF_RS04230 | 1.0286 | 2.30E-05 | HAD family haloacid dehalogenase hydrolase | Central metabolism |
| OG1RF_10836 | OG1RF_RS04385 | 1.0989 | 1.09E-07 | ABC superfamily ATP binding cassette transporter, membrane protein | Multifunction transporter |
| OG1RF_10838 | OG1RF_RS04395 | -1.8971 | 3.20E-12 | NRAMP family Mn2+/Fe2+ transporter (MntH2) | Multifunction transporter |
| OG1RF_10841 | OG1RF_RS04410 | 1.2338 | 1.77E-10 | peptide ABC transporter substrate-binding protein | Peptide ABC transporter |
| OG1RF_10842 | OG1RF_RS04415 | 1.0954 | 3.27E-08 | deaminase | Central metabolism |
| OG1RF_10843 | OG1RF_RS04420 | 1.2328 | 5.92E-09 | deaminase | Central metabolism |
| OG1RF_10849 | OG1RF_RS04450 | 1.0803 | 4.64E-07 | galK/galactokinase | Central metabolism |
| OG1RF_10869 | OG1RF_RS04550 | -5.8127 | 8.00E-16 | von Willebrand factor type A domain protein (ebpA) | Cell surface adhesin |
| OG1RF_10870 | OG1RF_RS04555 | -2.2992 | 6.51E-11 | cell wall surface anchor family protein (ebpB) | Cell surface adhesin |
| OG1RF_10871 | OG1RF_RS04560 | -2.2531 | 6.65E-11 | cell wall surface anchor family protein (ebpC) | Cell surface adhesin |
| OG1RF_10881 | OG1RF_RS04610 | 1.8894 | 2.58E-07 | hypothetical protein | Function unknown |
| OG1RF_10882 | OG1RF_RS04615 | 1.0794 | 5.82E-11 | amino acid permease | Peptide ABC transporter |
| OG1RF_10895 | OG1RF_RS04680 | 2.6415 | 2.57E-17 | amino acid ABC transporter permease | Peptide ABC transporter |
| OG1RF_10896 | OG1RF_RS04685 | 2.8651 | 2.32E-17 | glutamine ABC transporter permease | Peptide ABC transporter |
| OG1RF_10897 | OG1RF_RS04690 | 2.8198 | 2.85E-17 | glutamine ABC transporter substrate-binding protein | Peptide ABC transporter |
| OG1RF_10898 | OG1RF_RS04695 | 3.1924 | 1.63E-18 | ABC transporter ATP-binding protein glnQ | Multifunction transporter |
| OG1RF_10954 | OG1RF_RS04975 | -1.3335 | 5.66E-10 | luxS/S-ribosylhomocysteine lyase | Central metabolism |
| OG1RF_10990 | OG1RF_RS05150 | -3.4495 | 6.40E-15 | spermidine/putrescine ABC superfamily ATP binding cassette transporter, membrane protein | Peptide ABC transporter |
| OG1RF_10991 | OG1RF_RS05155 | -3.3045 | 3.27E-13 | permidine/putrescine ABC superfamily ATP binding cassette transporter, permease protein | Peptide ABC transporter |
| OG1RF_10992 | OG1RF_RS05160 | -3.3845 | 1.03E-14 | ABC superfamily ATP binding cassette | Peptide ABC transporter |
| OG1RF_10993 | OG1RF_RS05165 | -3.3332 | 3.06E-16 | spermidine/putrescine ABC superfamily ATP binding cassette transporter, spermidine/putrescine-binding | Peptide ABC transporter |
| OG1RF_10994 | OG1RF_RS05170 | -3.1835 | 5.56E-15 | ade/adenine deaminase | Central metabolism |
| OG1RF_10995 | OG1RF_RS05175 | -2.7434 | 4.48E-14 | mtaD/putative S-adenosylhomocysteine deaminase | Central metabolism |
| OG1RF_11000 | OG1RF_RS05200 | -2.3001 | 1.14E-10 | hypothetical protein | Function unknown |
| OG1RF_11001 | OG1RF_RS05205 | -2.2968 | 1.61E-12 | AP2 domain-containing protein | Function unknown |
| OG1RF_11031 | OG1RF_RS05360 | 1.6047 | 7.38E-07 | hypothetical protein | Function unknown |
| OG1RF_11032 | OG1RF_RS05365 | 1.5776 | 2.28E-08 | hypothetical protein | Function unknown |
| OG1RF_11036 | OG1RF_RS05385 | 1.0034 | 8.43E-12 | ATPase P | Respiration and ATP metabolism |
| OG1RF_11049 | OG1RF_RS05445 | -1.4193 | 4.83E-07 | DNA replication protein DnaD | DNA replication and repair |
| OG1RF_11050 | OG1RF_RS05450 | -1.4531 | 1.99E-09 | DNA replication protein DnaI | DNA replication and repair |
| OG1RF_11051 | OG1RF_RS05460 | -1.3359 | 6.83E-08 | hypothetical protein | Function unknown |
| OG1RF_11052 | OG1RF_RS05465 | -1.4973 | 1.40E-05 | RinA family transcriptional regulator | Transcription regulators |
| OG1RF_11054 | OG1RF_RS05470 | -2.0944 | 3.47E-13 | hypothetical protein | Function unknown |
| OG1RF_11055 | OG1RF_RS05475 | -2.0080 | 3.92E-13 | phage major tail protein, TP901-1 family | Cell surface adhesin |
| OG1RF_11056 | OG1RF_RS05480 | -1.9443 | 2.02E-12 | hypothetical protein | Function unknown |
| OG1RF_11057 | OG1RF_RS05485 | -2.0594 | 4.80E-12 | hypothetical protein | Function unknown |
| OG1RF_11058 | OG1RF_RS05490 | -2.1338 | 1.87E-14 | hypothetical protein | Function unknown |
| OG1RF_11059 | OG1RF_RS05495 | -2.2442 | 1.65E-13 | phage tail protein | Cell surface adhesin |
| OG1RF_11060 | OG1RF_RS05500 | -2.1279 | 2.17E-12 | phage structural protein | Cell surface adhesin |
| OG1RF_11061 | OG1RF_RS05505 | -2.0657 | 5.48E-12 | hypothetical protein | Function unknown |
| OG1RF_11062 | OG1RF_RS05510 | -2.0418 | 1.70E-11 | holin | Cell surface adhesin |
| OG1RF_11063 | OG1RF_RS05515 | -2.1428 | 4.62E-13 | endolysin | Cell surface adhesin |
| OG1RF_11070 | OG1RF_RS05550 | -0.6935 | 1.70E-06 | FtsW/RodA/SpovE family cell division protein | Central metabolism |
| OG1RF_11073 | OG1RF_RS05565 | 1.2714 | 4.28E-08 | LysR family transcriptional regulator | Transcription regulators |
| OG1RF_11134 | OG1RF_RS05880 | 1.0229 | 4.51E-07 | sugar ABC transporter permease | Multifunction transporter |
| OG1RF_11135 | OG1RF_RS05885 | 1.0375 | 5.68E-07 | sugar ABC superfamily ATP binding cassette transporter, membrane protein | Multifunction transporter |
| OG1RF_11157 | OG1RF_RS06005 | -1.2245 | 5.10E-05 | cro/CI family transcriptional regulator | Transcription regulators |
| OG1RF_11173 | OG1RF_RS06085 | 1.0652 | 4.00E-06 | molybdopterin-guanine dinucleotide biosynthesis protein A | Pyridine and purine metabolism |
| OG1RF_11175 | OG1RF_RS06095 | 1.1557 | 4.90E-04 | hypothetical protein | Function unknown |
| OG1RF_11178 | OG1RF_RS06110 | 1.2207 | 1.19E-08 | formate dehydrogenase subunit alpha | Central metabolism |
| OG1RF_11179 | OG1RF_RS06115 | 1.2463 | 1.84E-07 | molybdenum cofactor biosynthesis family protein | Central metabolism |
| OG1RF_11182 | OG1RF_RS06125 | 1.1060 | 2.90E-04 | GTP 3',8-cyclase MoaA | Respiration and ATP metabolism |
| OG1RF_11185 | OG1RF_RS06140 | 1.1033 | 4.90E-05 | molybdopterin-binding domain protein | Central metabolism |
| OG1RF_11227 | OG1RF_RS06355 | 4.7906 | 2.79E-16 | dipeptide epimerase | Central metabolism |
| OG1RF_11228 | OG1RF_RS06360 | 4.7032 | 1.08E-17 | transglutaminase domain protein | Central metabolism |
| OG1RF_11229 | OG1RF_RS06365 | 4.5861 | 1.32E-19 | peptide ABC transporter substrate-binding protein | Peptide ABC transporter |
| OG1RF_11239 | OG1RF_RS06415 | 1.6194 | 7.81E-12 | transglycosylase SLT domain protein | Central metabolism |
| OG1RF_11252 | OG1RF_RS06480 | 1.2736 | 6.68E-08 | hypothetical protein | Function unknown |
| OG1RF_11260 | OG1RF_RS06520 | 1.2215 | 1.45E-11 | brp/Blh family beta-carotene 15,15'-monooxygenase | Central metabolism |
| OG1RF_11294 | OG1RF_RS06685 | -1.1239 | 2.17E-12 | ABC superfamily ATP binding cassette transporter, ABC protein | Multifunction transporter |
| OG1RF_11295 | OG1RF_RS06690 | -1.0834 | 1.32E-12 | thyA/thymidylate synthase | Pyridine and purine metabolism |
| OG1RF_11306 | OG1RF_RS06745 | 1.2524 | 1.59E-10 | DNA mismatch repair protein MutT | DNA replication and repair |
| OG1RF_11308 | OG1RF_RS06755 | -1.5822 | 4.67E-08 | protease synthase | Protein synthesis, translation and degradation |
| OG1RF_11312 | OG1RF_RS06780 | -1.2487 | 2.90E-05 | DNA mismatch repair protein MutT | DNA replication and repair |
| OG1RF_11314 | OG1RF_RS06790 | -1.2863 | 6.60E-10 | katA/catalase | Central metabolism |
| OG1RF_11363 | OG1RF_RS07035 | 1.3264 | 2.52E-10 | CPBP family intramembrane metalloprotease | Central metabolism |
| OG1RF_11399 | OG1RF_RS07210 | 1.2029 | 6.60E-07 | hypothetical protein | Function unknown |
| OG1RF_11409 | OG1RF_RS07260 | 1.7773 | 6.72E-15 | NAD(P)H-dependent oxidoreductase | Respiration and ATP metabolism |
| OG1RF_11422 | OG1RF_RS07325 | -1.1815 | 2.77E-12 | carbonate dehydratase | Central metabolism |
| OG1RF_11423 | OG1RF_RS07330 | -1.1556 | 2.18E-12 | pyrE/orotate phosphoribosyltransferase | Protein synthesis, translation and degradation |
| OG1RF_11424 | OG1RF_RS07335 | -1.0443 | 1.91E-11 | pyrF/orotidine-5'-phosphate decarboxylase | Protein synthesis, translation and degradation |
| OG1RF_11425 | OG1RF_RS07340 | -1.1246 | 8.11E-14 | pyrDB/dihydroorotate oxidase | Protein synthesis, translation and degradation |
| OG1RF_11426 | OG1RF_RS07345 | -1.1162 | 1.19E-10 | pyrK/dihydroorotate dehydrogenase electron transfer subunit | Protein synthesis, translation and degradation |
| OG1RF_11427 | OG1RF_RS07350 | -1.1875 | 1.94E-12 | carB/carbamoyl-phosphate synthase, large subunit | Protein synthesis, translation and degradation |
| OG1RF_11428 | OG1RF_RS07355 | -1.5152 | 1.40E-11 | carA/carbamoyl-phosphate synthase, small subunit | Protein synthesis, translation and degradation |
| OG1RF_11429 | OG1RF_RS07360 | -1.7934 | 3.74E-13 | pyrC/dihydroorotase | Protein synthesis, translation and degradation |
| OG1RF_11430 | OG1RF_RS07365 | -1.9102 | 1.91E-15 | pyrB/aspartate carbamoyltransferase, catalytic subunit | Protein synthesis, translation and degradation |
| OG1RF_11431 | OG1RF_RS07370 | -2.1859 | 1.05E-15 | pyrP/NCS family uracil:cation symporter | Protein synthesis, translation and degradation |
| OG1RF_11432 | OG1RF_RS07375 | -2.3537 | 6.15E-14 | upp/uracil phosphoribosyltransferase | Pyridine and purine metabolism |
| OG1RF_11449 | OG1RF_RS07460 | -0.9800 | 5.02E-12 | tysrS2/tyrosine--tRNA ligase | Protein synthesis, translation and degradation |
| OG1RF_11451 | OG1RF_RS07470 | 3.0477 | 6.36E-13 | ABC transporter permease | Multifunction transporter |
| OG1RF_11452 | OG1RF_RS07475 | 1.9182 | 1.53E-08 | ccmA/ABC superfamily ATP binding cassette transporter, binding protein | Multifunction transporter |
| OG1RF_11480 | OG1RF_RS07605 | 1.0585 | 6.29E-07 | ABC superfamily ATP binding cassette transporter, ABC protein | Multifunction transporter |
| OG1RF_11501 | OG1RF_RS07710 | 1.3981 | 5.46E-10 | carbohydrate kinase | Central metabolism |
| OG1RF_11502 | OG1RF_RS07715 | -2.4254 | 3.63E-17 | oppA2/oligopeptide ABC superfamily ATP binding | Peptide ABC transporter |
| OG1RF_11503 | OG1RF_RS07720 | 2.2811 | 2.98E-11 | hypothetical protein | Function unknown |
| OG1RF_11534 | OG1RF_RS07875 | -1.6757 | 3.17E-09 | membrane protein | Cell surface adhesin |
| OG1RF_11535 | OG1RF_RS07880 | -1.8198 | 1.36E-08 | WxL domain surface protein | Cell surface adhesin |
| OG1RF_11536 | OG1RF_RS07885 | -2.0439 | 4.58E-09 | hypothetical protein | Function unknown |
| OG1RF_11537 | OG1RF_RS07890 | -1.5955 | 8.63E-12 | hypothetical protein | Function unknown |
| OG1RF_11670 | OG1RF_RS08560 | 1.4140 | 8.02E-10 | TetR family transcriptional regulator | Transcription regulators |
| OG1RF_11671 | OG1RF_RS08565 | 2.0753 | 1.00E-10 | hypothetical protein | Function unknown |
| OG1RF_11672 | OG1RF_RS08570 | 2.5207 | 2.86E-14 | major facilitator family transporter | Multifunction transporter |
| OG1RF_11678 | OG1RF_RS08605 | 1.1482 | 5.16E-08 | ABC superfamily ATP binding cassette transporter, membrane protein (efaB) | Multifunction transporter |
| OG1RF_11679 | OG1RF_RS08610 | 1.9362 | 1.12E-12 | ABC superfamily ATP binding cassette transporter, binding protein (efaA) | Multifunction transporter |
| OG1RF_11698 | OG1RF_RS08715 | -6.8130 | 1.52E-17 | cdaA/diadenylate cyclase | Central metabolism |
| OG1RF_11700 | OG1RF_RS08725 | -1.3000 | 5.26E-08 | hypothetical protein | Function unknown |
| OG1RF_11736 | OG1RF_RS08890 | 1.0178 | 2.12E-09 | group 2 glycosyl transferase | Central metabolism |
| OG1RF_11737 | OG1RF_RS08895 | 1.0102 | 4.00E-10 | group 2 glycosyl transferase | Central metabolism |
| OG1RF_11779 | OG1RF_RS09105 | 1.0066 | 3.39E-07 | hypothetical protein | Function unknown |
| OG1RF_11797 | OG1RF_RS09200 | -1.3334 | 9.37E-12 | pbuX/xanthine permease | Protein synthesis, translation and degradation |
| OG1RF_11798 | OG1RF_RS09205 | -1.7125 | 2.27E-12 | xpt/xanthine phosphoribosyltransferase | Protein synthesis, translation and degradation |
| OG1RF_11810 | OG1RF_RS09260 | -1.9969 | 3.04E-12 | amino acid permease | Protein synthesis, translation and degradation |
| OG1RF_11824 | OG1RF_RS09335 | 1.0078 | 2.90E-04 | hypothetical protein | Function unknown |
| OG1RF_11849 | OG1RF_RS09465 | 1.0036 | 5.32E-10 | zurR/Fur family transcriptional regulator ZurR | Transcription regulators |
| OG1RF_11860 | OG1RF_RS09520 | -1.3107 | 1.46E-12 | guaC/GMP reductase | Pyridine and purine metabolism |
| OG1RF_11861 | OG1RF_RS09525 | -1.6890 | 5.58E-10 | NCS2 family xanthine/uracil permease | Pyridine and purine metabolism |
| OG1RF_11862 | OG1RF_RS09530 | -1.7222 | 1.82E-11 | guaD/guanine deaminase | Pyridine and purine metabolism |
| OG1RF_11902 | OG1RF_RS09730 | -1.3517 | 4.13E-13 | argS/arginine--tRNA ligase | Protein synthesis, translation and degradation |
| OG1RF_11924 | OG1RF_RS09845 | -1.3779 | 8.35E-12 | cell wall surface anchor family protein | Cell surface adhesin |
| OG1RF_11929 | OG1RF_RS09875 | 2.0367 | 5.41E-14 | hypothetical protein | Function unknown |
| OG1RF_11938 | OG1RF_RS09915 | -2.8421 | 4.39E-18 | fumarate reductase | Respiration and ATP metabolism |
| OG1RF_11988 | OG1RF_RS10165 | -1.0762 | 5.57E-12 | atpA2/ATP synthase F1 sector alpha subunit | Respiration and ATP metabolism |
| OG1RF_11989 | OG1RF_RS10170 | -0.9976 | 7.53E-12 | atpH/ATP synthase F1 sector delta subunit | Respiration and ATP metabolism |
| OG1RF_11990 | OG1RF_RS10175 | -1.1357 | 7.57E-12 | atpF3/ATP synthase F0sector subunit B | Respiration and ATP metabolism |
| OG1RF_11991 | OG1RF_RS10180 | -1.1476 | 5.58E-08 | atpE2/ATP synthase F0 sector subunit C | Respiration and ATP metabolism |
| OG1RF_12013 | OG1RF_RS10300 | -4.1031 | 4.50E-19 | opuAA2/glycine betaine/L-proline ABC superfamily ATP binding cassette transporter, ABC protein | Peptide ABC transporter |
| OG1RF_12014 | OG1RF_RS10305 | -4.2795 | 7.57E-20 | glycine betaine/carnitine/choline ABC superfamily ATP binding cassette transporter, membrane/bindin | Peptide ABC transporter |
| OG1RF_12021 | OG1RF_RS10340 | -1.0354 | 4.68E-11 | potD/spermidine/putrescine ABC superfamily ATP binding cassette transporter, binding protein | Peptide ABC transporter |
| OG1RF_12022 | OG1RF_RS10345 | -1.0185 | 3.02E-09 | potC/spermidine/putrescine ABC superfamily ATP binding cassette transporter, membrane protein | Peptide ABC transporter |
| OG1RF_12023 | OG1RF_RS10350 | -1.0629 | 8.70E-11 | potB/spermidine/putrescine ABC superfamily ATP binding cassette transporter, membrane protein | Peptide ABC transporter |
| OG1RF_12024 | OG1RF_RS10355 | -0.9910 | 1.66E-10 | potA/spermidine/putrescine ABC superfamily ATP binding cassette transporter, ABC protein | Peptide ABC transporter |
| OG1RF_12031 | OG1RF_RS10390 | -1.5383 | 1.32E-10 | integral membrane protein | Cell surface adhesin |
| OG1RF_12037 | OG1RF_RS10420 | -1.0248 | 1.83E-10 | cutC/copper homeostasis protein CutC | Multifunction transporter |
| OG1RF_12041 | OG1RF_RS10440 | -1.1809 | 1.21E-11 | relA2/GTP diphosphokinase | Pyridine and purine metabolism |
| OG1RF_12094 | OG1RF_RS10710 | 1.2475 | 2.80E-06 | rpmG/50S ribosomal protein L33 | Protein synthesis, translation and degradation |
| OG1RF_12103 | OG1RF_RS10755 | -1.1747 | 7.69E-09 | ahpC/peroxiredoxin | Protein synthesis, translation and degradation |
| OG1RF_12117 | OG1RF_RS10830 | 1.1111 | 5.84E-10 | dinB/DNA-directed DNA polymerase IV | DNA replication and repair |
| OG1RF_12126 | OG1RF_RS10880 | -1.0452 | 2.30E-08 | recR/recombination protein R | DNA replication and repair |
| OG1RF_12131 | OG1RF_RS10905 | -1.0232 | 2.00E-05 | hypothetical protein | Function unknown |
| OG1RF_12132 | OG1RF_RS10910 | -1.0156 | 2.64E-09 | yniG/EmrB/QacA family drug resistance transporter | Multifunction transporter |
| OG1RF_12134 | OG1RF_RS10920 | 2.0022 | 1.70E-09 | MarR family transcriptional regulator | Transcription regulators |
| OG1RF_12144 | OG1RF_RS10970 | 1.4233 | 3.96E-13 | hypothetical protein | Function unknown |
| OG1RF_12145 | OG1RF_RS10975 | 1.0310 | 3.06E-10 | protein of hypothetical function DUF488 | Function unknown |
| OG1RF_12156 | OG1RF_RS11030 | -1.1243 | 2.29E-08 | hypothetical protein | Function unknown |
| OG1RF_12166 | OG1RF_RS11085 | 1.1749 | 7.51E-08 | hypothetical protein | Function unknown |
| OG1RF_12198 | OG1RF_RS11270 | 1.3071 | 3.60E-06 | hypothetical protein | Function unknown |
| OG1RF_12214 | OG1RF_RS11355 | -1.2334 | 3.32E-10 | greA/transcription elongation factor GreA | DNA replication and repair |
| OG1RF_12235 | OG1RF_RS11455 | 2.0615 | 1.94E-13 | S1 family extracellular protease | Protein synthesis, translation and degradation |
| OG1RF_12241 | OG1RF_RS11480 | 1.1324 | 3.70E-06 | LysR family transcriptional regulator | Transcription regulators |
| OG1RF_12244 | OG1RF_RS11495 | -1.7530 | 1.41E-09 | rbsK/ribokinase | Central metabolism |
| OG1RF_12277 | OG1RF_RS11660 | 1.2751 | 1.76E-07 | (S)-ureidoglycine aminohydrolase | Central metabolism |
| OG1RF_12278 | OG1RF_RS11665 | 1.9900 | 2.63E-09 | Zn-dependent hydrolase | Central metabolism |
| OG1RF_12279 | OG1RF_RS11670 | 2.4578 | 2.70E-13 | allB/allantoinase | Central metabolism |
| OG1RF_12280 | OG1RF_RS11675 | 2.3738 | 2.99E-12 | cytosine/purine permease | Pyridine and purine metabolism |
| OG1RF_12286 | OG1RF_RS11705 | 1.0431 | 5.12E-09 | hypothetical protein | Function unknown |
| OG1RF_12302 | OG1RF_RS11785 | -1.9781 | 5.37E-16 | DAACS family dicarboxylate/amino acid:cation symporter | Peptide ABC transporter |
| OG1RF_12311 | OG1RF_RS11840 | -3.4752 | 1.43E-16 | traC2/peptide ABC superfamily ATP binding cassette transporter, binding protein | Peptide ABC transporter |
| OG1RF_12322 | OG1RF_RS11895 | -1.0521 | 9.12E-09 | hypothetical protein | Function unknown |
| OG1RF_12328 | OG1RF_RS11925 | 1.1097 | 1.40E-06 | hypothetical protein | Function unknown |
| OG1RF_12332 | OG1RF_RS11945 | -0.9880 | 1.39E-08 | mreD/rod shape-determining protein MreD | Central metabolism |
| OG1RF_12376 | OG1RF_RS12175 | 1.1238 | 2.16E-08 | response regulator | Transcription regulators |
| OG1RF_12377 | OG1RF_RS12180 | 1.2229 | 5.48E-08 | sensor histidine kinase | Transcription regulators |
| OG1RF_12406 | OG1RF_RS12325 | 1.3831 | 1.14E-08 | RpiR family phosphosugar-binding transcriptional regulator | Transcription regulators |
| OG1RF_12407 | OG1RF_RS12330 | 1.4795 | 3.90E-10 | hypothetical protein | Function unknown |
| OG1RF_12451 | OG1RF_RS12525 | -1.1281 | 1.59E-08 | hypothetical protein | Function unknown |
| OG1RF_12452 | OG1RF_RS12530 | -1.1653 | 1.44E-07 | cell surface protein | Cell surface adhesin |
| OG1RF_12453 | OG1RF_RS12535 | -1.0167 | 2.05E-07 | cell surface protein | Cell surface adhesin |
| OG1RF_12454 | OG1RF_RS12540 | -1.0651 | 4.89E-07 | hypothetical protein | Function unknown |
| OG1RF_12456 | OG1RF_RS12550 | -1.0561 | 3.71E-11 | hypothetical protein | Function unknown |
| OG1RF_12460 | OG1RF_RS12565 | -1.1739 | 9.14E-10 | antiholin | Cell surface adhesin |
| OG1RF_12461 | OG1RF_RS12570 | -2.2240 | 1.11E-09 | lrgA/murein hydrolase regulator LrgA | Transcription regulators |
| OG1RF_12468 | OG1RF_RS12605 | 1.4241 | 3.65E-07 | rpsN2/30 S ribosomal protein S14 | Protein synthesis, translation and degradation |
| OG1RF_12469 | OG1RF_RS12610 | 1.6997 | 1.50E-04 | rpmG3/50S ribosomal protein L33 | Protein synthesis, translation and degradation |
| OG1RF_12472 | OG1RF_RS12625 | 1.3275 | 2.90E-06 | zinc ABC transporter substrate-binding protein AdcA | Multifunction transporter |
| OG1RF_12476 | OG1RF_RS12645 | -1.9367 | 3.81E-09 | PTS family fructose/mannitol (fru) porter component IIA | Central metabolism |
| OG1RF_12477 | OG1RF_RS12650 | -2.1041 | 3.59E-11 | PTS family ascorbate porter, IIB component | Central metabolism |
| OG1RF_12478 | OG1RF_RS12655 | -1.6053 | 1.28E-10 | PTS family fructose/mannitol (fru) porter component IIC | Central metabolism |
| OG1RF_12479 | OG1RF_RS12660 | -1.6381 | 2.56E-10 | PTS family porter component II | Central metabolism |
| OG1RF_12481 | OG1RF_RS12670 | -2.6669 | 8.30E-13 | choline binding protein | Central metabolism |
| OG1RF_12496 | OG1RF_RS12735 | 1.0877 | 7.72E-07 | DNA-binding protein | Transcription regulators |
| OG1RF_12499 | OG1RF_RS12745 | -1.5211 | 6.84E-14 | chitin binding protein | Central metabolism |
| OG1RF_12500 | OG1RF_RS12755 | 3.0987 | 4.16E-15 | LytR family transcriptional regulator | Transcription regulators |
| OG1RF_12501 | OG1RF_RS12760 | 1.5262 | 1.76E-08 | putative holin-like toxin | Cell surface adhesin |
| OG1RF_12508 | OG1RF_RS12790 | -1.1282 | 2.63E-09 | thiamine biosynthesis protein ApbE | Pyridine and purine metabolism |
| OG1RF_12509 | OG1RF_RS12795 | -1.5063 | 3.60E-11 | FMN-binding domain-containing protein | Respiration and ATP metabolism |
| OG1RF_12510 | OG1RF_RS12800 | -1.0783 | 2.27E-12 | ndh3/NADH dehydrogenase | Respiration and ATP metabolism |
| OG1RF_12520 | OG1RF_RS12850 | -1.5962 | 1.87E-12 | hypothetical protein | Function unknown |
| OG1RF_12521 | OG1RF_RS12855 | -1.4633 | 2.79E-14 | transcriptional regulator | Transcription regulators |
| OG1RF_12522 | OG1RF_RS12860 | -1.6410 | 1.52E-13 | hypothetical protein | Function unknown |
| OG1RF_12523 | OG1RF_RS12865 | -2.4802 | 1.58E-14 | hydantoinase/oxoprolinase | Central metabolism |
| OG1RF_12524 | OG1RF_RS12870 | -2.4324 | 8.19E-12 | hypothetical protein | Function unknown |
| OG1RF_12525 | OG1RF_RS12875 | -2.0522 | 3.64E-12 | cytosine/purine/uracil/thiamine/allantoin permease | Pyridine and purine metabolism |
| OG1RF_12560 | OG1RF_RS13105 | 1.5692 | 6.22E-11 | citXG/triphosphoribosyl-dephospho-CoA synthase | Central metabolism |
| OG1RF_12561 | OG1RF_RS13110 | 1.9845 | 1.91E-14 | oxaloacetate decarboxylase | Central metabolism |
| OG1RF_12562 | OG1RF_RS13115 | 1.9912 | 2.00E-15 | oadA/oxaloacetate decarboxylase | Central metabolism |
| OG1RF_12563 | OG1RF_RS13120 | 2.0634 | 2.28E-13 | citrate lyase holo-ACP synthase | Central metabolism |
| OG1RF_12564 | OG1RF_RS13125 | 2.0374 | 5.88E-14 | citrate lyase subunit alpha | Central metabolism |
| OG1RF_12565 | OG1RF_RS13130 | 2.2203 | 6.81E-13 | citrate lyase subunit beta | Central metabolism |
| OG1RF_12566 | OG1RF_RS13135 | 2.2249 | 7.22E-11 | citD/citrate lyase acyl carrier protein | Central metabolism |
| OG1RF_12567 | OG1RF_RS13140 | 2.4456 | 6.02E-16 | citC/citrate [pro-3S]-lyase] ligase | Central metabolism |
| OG1RF_12568 | OG1RF_RS13145 | 2.0942 | 4.64E-09 | hypothetical protein | Function unknown |
| OG1RF_12569 | OG1RF_RS13150 | 2.7291 | 4.15E-12 | gcdB/glutaconyl-CoA decarboxylase | Central metabolism |
| OG1RF_12570 | OG1RF_RS13155 | 2.4239 | 1.08E-11 | acetyl-CoA carboxylase biotin carboxyl carrier protein subunit | Central metabolism |
| OG1RF_12571 | OG1RF_RS13160 | 2.7063 | 1.37E-09 | hypothetical protein | Function unknown |
| OG1RF_12572 | OG1RF_RS13165 | 1.6313 | 6.21E-09 | citrate transporter | Multifunction transporter |
| OG1RF_10271 | OG1RF_RS13290 | 1.1906 | 8.32E-07 | acyl-CoA synthetase FdrA | Central metabolism |
| OG1RF_10825 | OG1RF_RS13410 | 1.3815 | 5.52E-10 | hypothetical protein | Function unknown |
| OG1RF_11048 | OG1RF_RS13435 | -1.4470 | 7.30E-05 | hypothetical protein | Function unknown |
| OG1RF_12429 | OG1RF_RS13650 | 1.3492 | 8.89E-09 | putative holin-like toxin | Cell surface adhesin |
