## Supplemental Table 2 for "c-di-AMP is essential for the virulence of *Enterococcus faecalis*"

**Table S2.** List of genes differentially expressed (FDR of 0.05 and 2-fold change cutoff) in *E. faecalis* ∆*dhhP∆gdpP* in comparison to OG1RF (wild type) strain.

| **Old Gene Locus** | **New Gene Locus** | **Log2FC** | **P-Value** | **Annotated Function** | **Functional Category** |
| --- | --- | --- | --- | --- | --- |
| OG1RF_10010 | OG1RF_RS00060 | -4.5015 | 1.54E-18 | DHH family protein/gdpP | Pyridine and purine metabolism |
| OG1RF_10011 | OG1RF_RS00065 | -2.0785 | 1.21E-14 | rplI/50S ribosomal protein L9 | Protein synthesis, translation and degradation |
| OG1RF_10028 | OG1RF_RS00145 | -1.2411 | 9.00E-08 | RidA family protein | Central metabolism |
| OG1RF_10030 | OG1RF_RS00155 | 1.1768 | 2.16E-04 | hypothetical protein | Function unknown |
| OG1RF_10058 | OG1RF_RS00295 | 1.4352 | 8.38E-05 | hypothetical protein | Function unknown |
| OG1RF_10059 | OG1RF_RS00300 | 1.0699 | 1.46E-06 | LLM class flavin-dependent oxidoreductase | Respiration and ATP metabolism |
| OG1RF_10083 | OG1RF_RS00415 | 1.1302 | 3.42E-09 | M protein trans-acting positive regulator | Transcriptional regulators |
| OG1RF_10091 | OG1RF_RS00455 | 1.4840 | 6.52E-13 | hypothetical protein | Function unknown |
| OG1RF_10092 | OG1RF_RS00460 | 3.2210 | 3.15E-12 | transcriptional regulator | Transcriptional regulators |
| OG1RF_10093 | OG1RF_RS00465 | 3.3365 | 4.45E-09 | L-serine dehydratase, iron-sulfur-dependent, beta subunit | Central metabolism |
| OG1RF_10094 | OG1RF_RS00470 | 2.8094 | 3.72E-11 | L-serine dehydratase, iron-sulfur-dependent,alpha subunit | Central metabolism |
| OG1RF_10095 | OG1RF_RS00475 | 3.0965 | 3.61E-14 | serine--tRNA ligase | Protein synthesis, translation and degradation |
| OG1RF_10098 | OG1RF_RS00490 | -0.9874 | 4.88E-11 | argR/arginine repressor | Central metabolism |
| OG1RF_10099 | OG1RF_RS00495 | -5.1092 | 6.81E-18 | arcA/arginine deiminase | Central metabolism |
| OG1RF_10100 | OG1RF_RS00500 | -5.6148 | 9.02E-17 | arcB/ornithine carbamoyltransferase | Central metabolism |
| OG1RF_10101 | OG1RF_RS00505 | -3.6044 | 1.12E-15 | arcC/carbamate kinase | Central metabolism |
| OG1RF_10102 | OG1RF_RS00510 | -2.3097 | 3.86E-14 | ntcA/global nitrogen regulator NtcA | Transcriptional regulators |
| OG1RF_10103 | OG1RF_RS00515 | -2.2901 | 2.74E-16 | C4-dicarboxylate ABC transporter | Peptide ABC transporter |
| OG1RF_10119 | OG1RF_RS00600 | 1.3986 | 6.38E-10 | adenosine deaminase | Central metabolism |
| OG1RF_10188 | OG1RF_RS00945 | 1.3561 | 7.33E-09 | N-acetyltransferase | Central metabolism |
| OG1RF_10198 | OG1RF_RS00990 | 1.1072 | 8.51E-07 | aldA/aldehyde dehydrogenase (NAD(+)) | Respiration and ATP metabolism |
| OG1RF_10199 | OG1RF_RS00995 | 1.4847 | 7.60E-14 | ldh2/L-lactate dehydrogenase | Respiration and ATP metabolism |
| OG1RF_10216 | OG1RF_RS01220 | -1.8713 | 3.27E-04 | bglA/6-phospho-beta-glucosidase | Central metabolism |
| OG1RF_10217 | OG1RF_RS01225 | -2.1098 | 1.08E-03 | phosphoglycerate mutase | Central metabolism |
| OG1RF_10288 | OG1RF_RS01590 | 1.2102 | 2.01E-11 | nhaC/Na+/H+ antiporter NhaC | Multifunction transporter |
| OG1RF_10304 | OG1RF_RS01670 | 1.1594 | 2.71E-05 | hypothetical protein | Function unknown |
| OG1RF_10305 | OG1RF_RS01675 | 1.2449 | 2.25E-08 | hypothetical protein | Function unknown |
| OG1RF_10388 | OG1RF_RS02095 | 1.1920 | 1.34E-10 | DeoR family transcriptional regulator | Transcriptional regulators |
| OG1RF_10389 | OG1RF_RS02100 | 1.1764 | 5.04E-12 | Xaa-Pro aminopeptidase" | Protein synthesis, translation and degradation |
| OG1RF_10455 | OG1RF_RS02420 | 1.0163 | 2.22E-09 | fruK2/1-phosphofructokinase | Central metabolism |
| OG1RF_10467 | OG1RF_RS02480 | 1.1715 | 4.79E-06 | putative thioredoxin | Respiration and ATP metabolism |
| OG1RF_10485 | OG1RF_RS02570 | -1.4649 | 2.06E-11 | cell wall surface anchor family protein | Cell surface adhesin |
| OG1RF_10486 | OG1RF_RS02575 | -1.3608 | 2.79E-11 | WxL domain surface protein | Cell surface adhesin |
| OG1RF_10487 | OG1RF_RS02580 | -1.3673 | 6.26E-11 | WxL domain surface protein | Cell surface adhesin |
| OG1RF_10488 | OG1RF_RS02585 | -1.4120 | 1.03E-13 | WxL domain surface protein | Cell surface adhesin |
| OG1RF_10489 | OG1RF_RS02590 | -1.3195 | 4.96E-10 | WxL domain surface protein | Cell surface adhesin |
| OG1RF_10490 | OG1RF_RS02595 | -1.0277 | 7.56E-10 | cell wall surface anchor family protein | Cell surface adhesin |
| OG1RF_10491 | OG1RF_RS02600 | -1.2613 | 4.96E-10 | hypothetical protein | Function unknown |
| OG1RF_10499 | OG1RF_RS02640 | 2.1280 | 1.41E-10 | hypothetical protein | Function unknown |
| OG1RF_10537 | OG1RF_RS02825 | 2.8184 | 3.30E-13 | aatB/amino acid ABC superfamily ATP binding cassette transporter, binding protein | Peptide ABC transporter |
| OG1RF_10538 | OG1RF_RS02830 | 2.4805 | 1.14E-13 | ABC superfamily ATP binding cassette transporter, ABC protein | Peptide ABC transporter |
| OG1RF_10539 | OG1RF_RS02835 | 1.8009 | 1.26E-09 | amino acid ABC superfamily ATP binding cassette transporter, membrane protein | Peptide ABC transporter |
| OG1RF_10580 | OG1RF_RS03040 | 1.1407 | 5.38E-11 | PemK family transcriptional regulator | Transcriptional regulators |
| OG1RF_10581 | OG1RF_RS03045 | 1.0104 | 7.35E-05 | hypothetical protein | Function unknown |
| OG1RF_10589 | OG1RF_RS03085 | 1.7993 | 3.78E-14 | cation efflux family protein (MntE) | Multifunction transporter |
| OG1RF_10602 | OG1RF_RS03145 | 1.1905 | 9.68E-06 | cro/CI family transcriptional regulator | Transcriptional regulators |
| OG1RF_10631 | OG1RF_RS03285 | 1.1823 | 1.34E-10 | mvk/mevalonate kinase | Central metabolism |
| OG1RF_10644 | OG1RF_RS03350 | 1.2299 | 4.59E-06 | hypothetical protein | Function unknown |
| OG1RF_10659 | OG1RF_RS03425 | 3.0407 | 6.84E-16 | hypothetical protein | Function unknown |
| OG1RF_10660 | OG1RF_RS03430 | 5.6652 | 1.23E-20 | rep/ATP-depenedent DNA helicase | DNA replication and repair |
| OG1RF_10664 | OG1RF_RS03450 | 1.1553 | 1.10E-08 | hypothetical protein | Function unknown |
| OG1RF_10679 | OG1RF_RS03525 | 1.8784 | 1.38E-09 | hypothetical protein | Function unknown |
| OG1RF_10745 | OG1RF_RS03850 | -1.8982 | 5.76E-08 | PTS sugar transporter subunit IIB | Central metabolism |
| OG1RF_10750 | OG1RF_RS03875 | -2.1261 | 8.76E-10 | PTS sugar transporter subunit IIB | Central metabolism |
| OG1RF_10751 | OG1RF_RS03880 | -1.4775 | 6.21E-08 | PTS mannose transporter subunit IIA | Central metabolism |
| OG1RF_10766 | OG1RF_RS03960 | 2.1415 | 4.30E-16 | 6-aminohexanoate hydrolase | Central metabolism |
| OG1RF_10805 | OG1RF_RS04230 | 1.0664 | 6.56E-06 | HAD family haloacid dehalogenase hydrolase | Central metabolism |
| OG1RF_10807 | OG1RF_RS04240 | 1.0510 | 7.64E-04 | glutathione S-transferase | Central metabolism |
| OG1RF_10838 | OG1RF_RS04395 | -1.7405 | 2.09E-12 | NRAMP family Mn2+/Fe2+ transporter (MntH2) | Multifunction transporter |
| OG1RF_10875 | OG1RF_RS04580 | -1.1436 | 3.54E-11 | hypothetical protein | Function unknown |
| OG1RF_10876 | OG1RF_RS04585 | -1.0540 | 1.44E-09 | hypothetical protein | Function unknown |
| OG1RF_10881 | OG1RF_RS04610 | 1.3812 | 8.07E-06 | hypothetical protein | Function unknown |
| OG1RF_10959 | OG1RF_RS05000 | -1.1472 | 1.06E-03 | hypothetical protein | Function unknown |
| OG1RF_10985 | OG1RF_RS05125 | 3.4665 | 1.21E-18 | alsS/acetolactate synthase | Central metabolism |
| OG1RF_10986 | OG1RF_RS05130 | 3.7926 | 4.52E-19 | budA/alpha-acetolactate decarboxylase | Central metabolism |
| OG1RF_10994 | OG1RF_RS05170 | -1.2519 | 6.59E-11 | ade/adenine deaminase | Central metabolism |
| OG1RF_10995 | OG1RF_RS05175 | -1.4783 | 1.91E-11 | mtaD/putative S-adenosylhomocysteine deaminase | Central metabolism |
| OG1RF_11002 | OG1RF_RS05215 | 1.1028 | 6.50E-06 | hypothetical protein | Function unknown |
| OG1RF_11019 | OG1RF_RS05300 | -1.0406 | 1.01E-04 | hypothetical protein | Function unknown |
| OG1RF_11027 | OG1RF_RS05340 | -1.0092 | 2.62E-04 | hypothetical protein | Function unknown |
| OG1RF_11074 | OG1RF_RS05570 | -1.1023 | 2.78E-10 | mgtA/magnesium-importing ATPase | Multifunction transporter |
| OG1RF_11095 | OG1RF_RS05675 | -1.0086 | 7.24E-07 | hypothetical protein | Function unknown |
| OG1RF_11102 | OG1RF_RS05720 | 1.1677 | 1.60E-03 | hypothetical protein | Function unknown |
| OG1RF_11130 | OG1RF_RS05860 | 1.1026 | 2.69E-10 | peptidase M4 | Protein synthesis, translation and degradation |
| OG1RF_11141 | OG1RF_RS05920 | 2.0138 | 8.19E-14 | pdhA/pyruvate dehydrogenase complex E1 component alpha subunit | Respiration and ATP metabolism |
| OG1RF_11142 | OG1RF_RS05925 | 2.1330 | 6.63E-15 | pdhB/pyruvate dehydrogenase complex E1 component beta subunit | Respiration and ATP metabolism |
| OG1RF_11143 | OG1RF_RS05930 | 2.3812 | 7.80E-15 | aceF/pyruvate dehydrogenase complex E2, dihydrolipoamide acetyltransferase | Respiration and ATP metabolism |
| OG1RF_11144 | OG1RF_RS05935 | 2.5945 | 7.66E-15 | lpd/dihydrolipoyl dehydrogenase | Central metabolism |
| OG1RF_11145 | OG1RF_RS05940 | 1.0187 | 1.73E-08 | AraC family transcriptional regulator | Transcriptional regulators |
| OG1RF_11159 | OG1RF_RS06015 | 1.3422 | 3.87E-11 | metal-dependent hydrolase | Central metabolism |
| OG1RF_11160 | OG1RF_RS06020 | 1.3739 | 2.79E-13 | hypothetical protein | Function unknown |
| OG1RF_11161 | OG1RF_RS06025 | -5.0299 | 1.40E-17 | dhhP/DHH family protein | Pyridine and purine metabolism |
| OG1RF_11165 | OG1RF_RS06045 | -0.9925 | 1.15E-06 | DEAD/DEAH box family ATP-dependent RNA helicase | DNA replication and repair |
| OG1RF_11225 | OG1RF_RS06345 | 1.8757 | 1.62E-13 | N-acetyltransferase | Central metabolism |
| OG1RF_11227 | OG1RF_RS06355 | 1.2646 | 9.37E-08 | dipeptide epimerase | Central metabolism |
| OG1RF_11228 | OG1RF_RS06360 | 1.1969 | 7.21E-09 | transglutaminase domain protein | Central metabolism |
| OG1RF_11235 | OG1RF_RS06395 | -1.3387 | 4.60E-04 | PTS fructose transporter subunit IIA | Central metabolism |
| OG1RF_11260 | OG1RF_RS06520 | 1.0458 | 8.21E-11 | brp/Blh family beta-carotene 15,15'-monooxygenase | Central metabolism |
| OG1RF_11301 | OG1RF_RS06720 | 1.3311 | 1.29E-12 | MFS transporter | Multifunction transporter |
| OG1RF_11308 | OG1RF_RS06755 | 1.6223 | 2.14E-10 | protease synthase | Central metabolism |
| OG1RF_11313 | OG1RF_RS06785 | -1.1117 | 5.79E-08 | hypothetical protein | Function unknown |
| OG1RF_11421 | OG1RF_RS07320 | -1.0867 | 9.55E-12 | LysR family transcriptional regulator | Transcriptional regulators |
| OG1RF_11422 | OG1RF_RS07325 | -1.2801 | 4.91E-13 | cah/carbonate dehydratase | Pyridine and purine metabolism |
| OG1RF_11423 | OG1RF_RS07330 | -1.3189 | 2.75E-13 | pyrE/orotate phosphoribosyltransferase | Pyridine and purine metabolism |
| OG1RF_11424 | OG1RF_RS07335 | -1.3245 | 5.31E-13 | pyrF/orotidine-5'-phosphate decarboxylase | Pyridine and purine metabolism |
| OG1RF_11425 | OG1RF_RS07340 | -1.2331 | 1.96E-14 | pyrDB/dihydroorotate oxidase | Pyridine and purine metabolism |
| OG1RF_11426 | OG1RF_RS07345 | -1.2655 | 1.84E-11 | pyrK/dihydroorotate dehydrogenase electron transfer subunit | Pyridine and purine metabolism |
| OG1RF_11427 | OG1RF_RS07350 | -1.2078 | 1.83E-12 | pyrK/dihydroorotate dehydrogenase electron transfer subunit | Pyridine and purine metabolism |
| OG1RF_11428 | OG1RF_RS07355 | -1.1894 | 3.65E-10 | carA/carbamoyl-phosphate synthase, small subunit | Pyridine and purine metabolism |
| OG1RF_11429 | OG1RF_RS07360 | -1.1728 | 1.50E-10 | pyrC/dihydroorotase | Pyridine and purine metabolism |
| OG1RF_11430 | OG1RF_RS07365 | -1.0975 | 3.02E-12 | pyrB/aspartate carbamoyltransferase, catalytic subunit | Pyridine and purine metabolism |
| OG1RF_11431 | OG1RF_RS07370 | -1.0076 | 2.72E-11 | pyrP/NCS family uracil:cation symporter | Multifunction transporter |
| OG1RF_11442 | OG1RF_RS07425 | 1.0865 | 9.23E-13 | rnhA/ribonuclease HI | DNA replication and repair |
| OG1RF_11524 | OG1RF_RS07825 | 1.0018 | 7.43E-07 | hypothetical protein | Function unknown |
| OG1RF_11533 | OG1RF_RS07870 | -1.1466 | 1.82E-09 | zinc-dependent alcohol dehydrogenase | Central metabolism |
| OG1RF_11534 | OG1RF_RS07875 | -1.6529 | 5.58E-10 | cell surface protein | Cell surface adhesin |
| OG1RF_11535 | OG1RF_RS07880 | -1.8928 | 1.36E-09 | hypothetical protein | Function unknown |
| OG1RF_11536 | OG1RF_RS07885 | -1.6583 | 5.06E-09 | hypothetical protein | Function unknown |
| OG1RF_11537 | OG1RF_RS07890 | -1.6052 | 1.58E-12 | hypothetical protein | Function unknown |
| OG1RF_11670 | OG1RF_RS08560 | 1.0173 | 4.41E-08 | TetR family transcriptional regulator | Transcriptional regulators |
| OG1RF_11685 | OG1RF_RS08645 | 1.0389 | 1.50E-03 | hypothetical protein | Function unknown |
| OG1RF_11753 | OG1RF_RS08975 | 1.2938 | 2.40E-10 | PTS maltose transporter subunit IIBC | Central metabolism |
| OG1RF_11799 | OG1RF_RS09210 | 1.2386 | 2.86E-11 | hydrolase | Central metabolism |
| OG1RF_11813 | OG1RF_RS09275 | 1.1412 | 1.85E-03 | hypothetical protein | Function unknown |
| OG1RF_11927 | OG1RF_RS09860 | 1.0218 | 9.98E-07 | azlC/LIV-E family branched chain amino acid exporter AzlC | Peptide ABC transporter |
| OG1RF_11928 | OG1RF_RS09865 | 1.0319 | 4.62E-06 | azlD/LIV-E family branched chain amino acid exporter AzlD | Peptide ABC transporter |
| OG1RF_11939 | OG1RF_RS09920 | 1.1697 | 8.17E-13 | Ktr system potassium uptake protein D | Multifunction transporter |
| OG1RF_12013 | OG1RF_RS10300 | 1.3323 | 5.32E-12 | glycine/betaine ABC transporter ATP-binding protein | Peptide ABC transporter |
| OG1RF_12014 | OG1RF_RS10305 | 1.0652 | 3.21E-11 | glycine/betaine ABC transporter permease | Peptide ABC transporter |
| OG1RF_12071 | OG1RF_RS10595 | -0.9875 | 9.44E-10 | transcriptional regulator | Transcriptional regulators |
| OG1RF_12127 | OG1RF_RS10885 | 1.0053 | 9.43E-08 | tenA/thiaminase II | Central metabolism |
| OG1RF_12134 | OG1RF_RS10920 | 1.5463 | 3.97E-08 | MarR family transcriptional regulator | Transcriptional regulators |
| OG1RF_12135 | OG1RF_RS10925 | 1.3935 | 2.20E-11 | thiD2/phosphomethylpyrimidine kinase | Pyridine and purine metabolism |
| OG1RF_12136 | OG1RF_RS10930 | 1.2765 | 6.21E-10 | thiE/thiamine-phosphate diphosphorylase | Pyridine and purine metabolism |
| OG1RF_12137 | OG1RF_RS10935 | 1.1645 | 1.34E-08 | thiM/putative hydroxyethylthiazole kinase | Pyridine and purine metabolism |
| OG1RF_12138 | OG1RF_RS10940 | 1.1580 | 1.14E-07 | energy coupling factor transporter S component ThiW | Respiration and ATP metabolism |
| OG1RF_12155 | OG1RF_RS11025 | 1.5728 | 1.11E-11 | brp/Blh family beta-carotene 15,15'-monooxygenase | Central metabolism |
| OG1RF_12156 | OG1RF_RS11030 | -0.9896 | 5.01E-08 | hypothetical protein | Function unknown |
| OG1RF_12193 | OG1RF_RS11220 | 1.2226 | 8.14E-12 | group 1 glycosyl transferase | Central metabolism |
| OG1RF_12229 | OG1RF_RS11425 | 1.0622 | 1.10E-11 | hypothetical protein | Function unknown |
| OG1RF_12235 | OG1RF_RS11455 | -1.0052 | 1.98E-08 | S1 family extracellular protease | Cell surface adhesin |
| OG1RF_12244 | OG1RF_RS11495 | -1.4849 | 1.15E-09 | rbsK/ribokinase | Protein synthesis, translation and degradation |
| OG1RF_12295 | OG1RF_RS11750 | 1.3242 | 1.68E-13 | serine/threonine transporter SstT | Multifunction transporter |
| OG1RF_12298 | OG1RF_RS11765 | 1.8377 | 5.41E-15 | hypothetical protein | Function unknown |
| OG1RF_12299 | OG1RF_RS11770 | 6.2948 | 1.97E-20 | hypothetical protein | Function unknown |
| OG1RF_12300 | OG1RF_RS11775 | 6.2301 | 4.68E-18 | hypothetical protein | Function unknown |
| OG1RF_12376 | OG1RF_RS12175 | 1.1352 | 8.82E-09 | response regulator | Transcriptional regulators |
| OG1RF_12377 | OG1RF_RS12180 | 1.0284 | 2.86E-07 | sensor histidine kinase | Transcriptional regulators |
| OG1RF_12406 | OG1RF_RS12325 | 1.2259 | 3.48E-08 | MurR/RpiR family transcriptional regulator | Transcriptional regulators |
| OG1RF_12425 | OG1RF_RS12410 | 1.6214 | 4.95E-13 | glycosyl hydrolase | Central metabolism |
| OG1RF_12426 | OG1RF_RS12415 | 1.6538 | 1.82E-12 | yudM/beta-phosphoglucomutase | Central metabolism |
| OG1RF_12449 | OG1RF_RS12515 | 1.0327 | 9.07E-10 | M protein trans-acting positive regulator | Transcriptional regulators |
| OG1RF_12450 | OG1RF_RS12520 | 1.1504 | 1.25E-10 | hypothetical protein | Function unknown |
| OG1RF_12451 | OG1RF_RS12525 | 1.5783 | 2.68E-12 | hypothetical protein | Function unknown |
| OG1RF_12452 | OG1RF_RS12530 | 1.9879 | 1.35E-12 | cell surface protein | Cell surface adhesin |
| OG1RF_12453 | OG1RF_RS12535 | 1.9102 | 8.19E-13 | cell surface protein | Cell surface adhesin |
| OG1RF_12454 | OG1RF_RS12540 | 1.9745 | 1.75E-12 | hypothetical protein | Function unknown |
| OG1RF_12455 | OG1RF_RS12545 | 2.0156 | 7.39E-12 | hypothetical protein | Function unknown |
| OG1RF_12456 | OG1RF_RS12550 | 1.5895 | 1.78E-14 | hypothetical protein | Function unknown |
| OG1RF_12460 | OG1RF_RS12565 | -1.3021 | 5.86E-11 | antiholin | Cell surface adhesin |
| OG1RF_12461 | OG1RF_RS12570 | -2.2234 | 1.69E-10 | lrgA/murein hydrolase regulator LrgA | Transcriptional regulators |
| OG1RF_12464 | OG1RF_RS12585 | 1.2533 | 3.70E-11 | ABC superfamily ATP binding cassette transporter, binding protein (Fe) | Multifunction transporter |
| OG1RF_12465 | OG1RF_RS12590 | 1.1055 | 5.08E-09 | metal ion ABC superfamily ATP binding cassette transporter, membrane protein (Fe) | Multifunction transporter |
| OG1RF_12466 | OG1RF_RS12595 | 1.3893 | 2.99E-10 | ABC superfamily ATP binding cassette transporter, ABC protein (Fe) | Multifunction transporter |
| OG1RF_12476 | OG1RF_RS12645 | -2.0029 | 3.70E-10 | PTS mannose transporter subunit IIA | Central metabolism |
| OG1RF_12477 | OG1RF_RS12650 | -2.1065 | 4.51E-12 | PTS sorbose transporter subunit IIB | Central metabolism |
| OG1RF_12478 | OG1RF_RS12655 | -1.9464 | 2.81E-12 | PTS mannose transporter subunit IIC | Central metabolism |
| OG1RF_12479 | OG1RF_RS12660 | -1.6888 | 2.68E-11 | PTS mannose transporter subunit IID | Central metabolism |
| OG1RF_12499 | OG1RF_RS12745 | -1.2299 | 3.91E-13 | chitin binding protein | Central metabolism |
| OG1RF_12538 | OG1RF_RS12995 | 1.0213 | 1.19E-11 | IMP dehydrogenase | Central metabolism |
| OG1RF_11021 | OG1RF_RS13430 | 1.4102 | 2.74E-05 | hypothetical protein | Function unknown |
| OG1RF_11716 | OG1RF_RS13525 | 1.0237 | 2.05E-05 | hypothetical protein | Function unknown |
| OG1RF_12407 | OG1RF_RS13630 | 1.6431 | 4.19E-11 | hypothetical protein | Function unknown |
