## Supplemental Table 3 for "c-di-AMP is essential for the virulence of *Enterococcus faecalis*"

**Table S3.** Primers used in this study.

|  | Primer name | Primer sequence (5’ 🡪 3’) | Restriction sites^a^ |
| --- | --- | --- | --- |
| Primers for qRT-PCR | ebpA_F’ | GTGGGAGCCTTTGAATTG | - |
|  | ebpA_R’ | CTCATGTCCTGCAGGTGC | - |
|  | cdaA_F’ | GCCTGAATAATACGCTCATCTTCTTGT | - |
|  | cdaA_R’ | GAACAAAAGCAGTCCAACTATTAAAAGG | - |
|  | gdpP_F’ | AGACACAATGCCAATAGCCGTTTG | - |
|  | gdpP_R’ | GCCAAGTTACGTCTTATTAGAATTAGCAG | - |
|  | dhhP_F’ | TGAACCCTATGGAGATTTAGTTTGG | - |
|  | dhhP_R’ | TGGTCTAACTCACGATTTAAGTCTGC | - |
| Primers for cloning in pCJK47 vector | cdaA_Arm1 F’ | GACTGG**GGATCC**TTGGTTGTA | BamHI |
|  | cdaA_Arm1 R’ | CGTAAGGAGT**GAATTC**CGGAATCATC | EcoRI |
|  | cdaA_Arm2 F’ | GATTCA**GAATTC**ACGACATCATACCCC | EcoRI |
|  | cdaA_Arm2 R’ | GGAGAC**GCATGC**GCTTACG | SphI |
|  | dhhP_Arm1 F’ | GTGAAG**CTGCAG**CGTATCG | PstI |
|  | dhhP_Arm1 R’ | GCTGCCAT**GAATTC**TTTTACTACG | EcoRI |
|  | dhhP_Arm2 F’ | GCTTGCCAA**GAATTC**GCGCAAAAATAA | EcoRI |
|  | dhhP_Arm2 R’ | CAGTTTCGTTTAAAAT**CCCGGG**AATAA | XmaI |
|  | gdpP_Arm1 F’ | GATCCT**GGATCC**CGTTACACCG | BamHI |
|  | gdpP_Arm1 R’ | CGTTTTTTTGAATT**CCCGGG**TTTTGCAT | XmaI |
|  | gdpP_Arm2 F’ | GTATAA**CCCGGG**AACACGCGC | XmaI |
|  | gdpP_Arm2 R’ | GCGCTC**GCATGC**TTCTGG | SphI |
|  | pcJK47 F’ | GCATGTTGATACGCTTG | - |
|  | pcJK47 R’ | AACATGTATTCACGAACGAA | - |
| Primers for screening deletion mutants | gdpP_F’ | GAAAGATTGGAGGCAAAAAAATGC | - |
|  | gdpP_R’ | GACTTTCATTTTCTTCACTCCTGTTC | - |
|  | cdaA_F’ | GCAATGAAAGCAAACCATTAAACCAGC | - |
|  | cdaA_R’ | GCGTTCTGTGATGAAAGAAAATAATGTGG | - |
|  | dhhP_F’ | CGTCGCTCTGCAAAATTAGATATTGAAG | - |
|  | dhhP_R’ | GTAGTCCTAATGATTTATCAACGTCCGATAAC | - |
|  | pheS F’ | GGATGGTAACACGATAGCTCCTTCC |  |
|  | pheS R’ | TTCAGGTACAGCAACGCCCTCAAC |  |
| Primers for cloning in pCIE vector | pCIE F’ | TTGTATAAATGTTGGAGCAGCG | - |
|  | pCIE R’ | ATCGTGTTTTTCTTGGAATTGTGC | - |
|  | cdaA F’ | CTTG**GGATCC**AAGAGGAGGTGAGGGGTATGATGTCGTTTCAATT | BamHI |
|  | cdaA R’ | GCTTTTTT**GCATGC**CTTTTCTCATTTG | SphI |
|  | pde F’ | GAAC**GGATCC**GGCAGGAGGTACAAAGAATGGACGTAGTAAAAGAAATTATGG | BamHI |
|  | pde R’ | CTACTT**GCATGC**TACTTAATTTTTGCG | SphI |
|  | gdpp F’ | CAGAA**GGATCC**GAGAGGAGGGCAAAAAAATGCAAAAGAGAATTCA | BamHI |
|  | gdpp R’ | GACTTT**GCATGC**CTTCACTC | SphI |

^a^ Restriction sites are underlined and bold in the primer sequence.
